# MSstatsBioNet: Integrating Statistical Analyses with Prior Knowledge Biomolecular Networks for Quantitative Proteomics and Phosphoproteomics

**DOI:** 10.64898/2026.07.09.737605

**Authors:** Anthony Wu, Devon Kohler, Pruthvi Prakash Navada, Julia E. Robbins, Gabriel E. Boyle, Alex Boshart, Klas Karis, Jacques Neefjes, Ana Konvalinka, Jay Sarthy, Lindsay Pino, Benjamin M. Gyori, Olga Vitek

## Abstract

A common outcome of quantitative mass spectrometry-based proteomic and phosphoproteomic experiments is a list of proteins that are differentially abundant between conditions. However, biological interpretation requires evaluation in the context of prior knowledge of biological mechanisms and protein function. One approach to facilitate mechanistic biological interpretation is to integrate such lists with biological network databases, built from manually curated resources and text mining systems. This manuscript automates this process with MSstatsBioNet, a Bioconductor package that integrates MSstats, a family of open-source packages for detecting differentially abundant proteins, and INDRA, a system that extracts biomolecular networks from biomedical literature using text mining and merges those networks with the content of curated knowledge bases. Taking as input a list of differentially abundant proteins from MSstats, MSstatsBioNet retrieves a protein subnetwork from INDRA and overlays experimental fold changes onto the underlying subnetwork. Users can then interact with the network and overlaid data, interrogating primary literature evidence to construct granular mechanistic narratives for iterative hypothesis generation. We demonstrate the utility of this approach with three case studies, two measuring changes in protein abundance and one measuring changes in phosphorylation.

## Introduction

Bottom-up liquid chromatography and mass spectrometry (LC-MS)-based proteomics is an indispensable tool for studying protein abundances,^1,2^ post-translational modifications (PTMs),^3,4^ and interactions.^5,6^ The predominant data acquisition modes are selected reaction monitoring (SRM)^7^ and parallel reaction monitoring (PRM)^8^ for targeted workflows, and data-dependent acquisition (DDA)^9^ and data-independent acquisition (DIA)^10^ for discovery workflows. Labeling methods, such as tandem mass tags (TMT),^11^ aim to improve sample throughput by combining multiple samples into a single mixture.

A key step in the analysis of quantitative proteomic experiments with all the workflows is the detection of proteins that are differentially abundant between conditions. Numerous software packages have been developed for this task, including MSstats,^12,13^ MSqRob2,^14–16^ DEqMS,^17,18^ prolfqua,^19^ proDA,^20^ and limpa.^21^ Some packages develop broader software ecosystems that handle various aspects of the workflow. For example, the MSstats ecosystem of R/Bioconductor packages converts the output of various spectral processing tools that identify and quantify the abundance of analytes into a unified MSstats-compatible format. ^13^ The MSstats ecosystem also adapts to different workflows, such as TMT-based workflows with MSstatsTMT^22^ and PTM-based workflows with MSstatsPTM.^23^

Although many tools for differential analysis exist, interpreting the results of a differential analysis remains challenging. ^24^ To facilitate interpretation, experimentalists rely on prior knowledge, defined here as existing curated and literature-derived information about biological pathways, protein function, perturbation targets, and disease mechanisms. By integrating differential analyses with prior knowledge, biological interpretation can focus on a tractable set of plausible explanations.

Integration of differential analysis with prior knowledge can take multiple forms. One common approach is functional enrichment analysis, which compares a set of differentially abundant genes or proteins with a set of possible biological processes, molecular functions, or pathways. Two popular methods are over-representation analysis (ORA),^25^ which is implemented in DAVID,^26^ clusterProfiler,^27^ limma,^28^ and GOseq,^29^ and gene set enrichment analysis (GSEA),^30^ which is implemented in fgsea^31^ and clusterProfiler,^27^ with a hallmark list of gene sets established in MSigDB.^32,33^ At the level of PTMs, kinase-substrate enrichment analysis tools such as KEA3^34^ infer whether certain upstream kinases are over-represented among a list of substrates.

Despite the broad availability of methods and implementations, functional enrichment analyses are usually insufficient at generating testable hypotheses, i.e. causal chains of mechanistic connection between an experimental perturbation and a subset of observed protein changes, rooted in prior literature and verifiable via a follow-up experiment.

Databases of biological networks support an alternative approach for building mechanistic narratives for hypothesis generation. The databases can be loosely grouped into those that contain manually curated relationships collected from literature or data, and those that establish these relationships by automated mining of text and data. Manually curated databases offer reliable mechanistic knowledge vetted by human experts. They can be categorized into domain-agnostic databases, such as Omnipath,^35^ Reactome,^36^ Pathway Commons,^37^ PhosphoSitePlus,^38^ and domain-specific databases, such as NephroDIP.^39^ However, due to the manual nature of curation, most cover only a fraction of the available literature and have limited scalability and coverage.

Alternatively, text mining systems extract biological networks from a large corpora of literature, providing broad coverage that is advantageous for building mechanistic narratives. Text mining systems for biomedical literature include Reach, ^40^ Sparser,^41^ and RLIMS-P.^42^ In contrast to standalone text mining systems, knowledge assemblers aggregate prior knowledge using multiple text mining systems into a single knowledge base. Examples of knowledge assemblers include STRING^43,44^ and INDRA.^45^ STRING was originally designed to aggregate protein-protein interactions and functional associations from literature. In contrast to STRING, INDRA is more mechanistic in nature as it aggregates networks such as signaling pathways and protein complexes, as well as PTM site-specific mechanisms.

Several tools integrate statistical analysis results, such as correlations between proteins or lists of differentially abundant proteins, with biological network databases for the purpose of network visualization and exploration. However, these tools share several limitations that hinder iterative hypothesis generation. First, tools such as ChiBE,^46^ CausalPath,^47^ Pathway Pilot,^48^ Pathway Volcano,^49^ ReactomeFIViz^50^ and PTMNavigator^51^ rely exclusively on manually curated databases, which, as mentioned above, cover only a fraction of the available literature and are therefore insufficient for building mechanistic narratives. In contrast to querying manually curated databases, SPONGE^52^ and OmicsVisualizer^53^ query STRING. However, SPONGE is limited to using STRING for interpretation of potential protein-protein interactions while OmicsVisualizer only supports interpretation of results in the context of protein associations in literature, which does not fully capture mechanistic behavior. The second limitation of these tools is the lack of a streamlined interface to directly inspect the individual pieces of evidence underlying each statement or to flag incorrect extractions. As a result, users are unable to assess or improve the reliability of the retrieved network. Finally, none of the tools provides flexible filtering by both experimental parameters, such as log_2_ fold change and adjusted p-value, and by properties of the retrieved network statements themselves, such as the amount of supporting evidence and the type of biological relationship, all of which are necessary for iteratively narrowing a large network to a tractable set of hypotheses.

To address these limitations, we propose a workflow for iterative generation of hypotheses underlying the results of quantitative mass spectrometry-based proteomic experiments for human samples. The workflow integrates a list of differentially abundant proteins or phosphosites with the information in the INDRA knowledge base structured as a mechanistic biomolecular network. The workflow overcomes the limitations described above by querying mechanistic relationships from a broad coverage of literature, extracting PTM-site specific mechanisms for phosphoproteomics experiments, providing a streamlined interface for inspecting and reviewing the evidence behind extracted relationships, and offering a variety of filters for quick and iterative querying which allows rapid hypothesis generation. The workflow is implemented in MSstatsBioNet, a freely available open-source R package compatible with the MSstats family of statistical software packages, and is available at Bioconductor. MSstatsBioNet is also accessible via the MSstatsShiny graphical user interface.^54^ Below we present details of the approach, as well as three case studies demonstrating its utility for interpreting experimental results and forming novel hypotheses for future experimental validation and application.

## Background

### Experimental design and spectral processing

**Figure 1** summarizes the full MS proteomics experiment workflow, from experimental design and data acquisition through the MSstats ecosystem to MSstatsBioNet’s subnetwork extraction and visualization steps. A study starts with an experimental design (step 1 of **Figure 1**) to confirm or reject a hypothesis. An experimental design consists of conditions of interest, such as presence or absence of a disease, or exposure to a chemical perturbation.^55^ The design must specify the biological subjects, such as cell lines, patients, or animals, representing each condition. The design also defines other technical factors that provide additional context to the data to be acquired, such as enrichment protocols, data acquisition mode, and instrument type, as well as dose amount and duration in the case of chemical perturbations.

**Figure 1:**
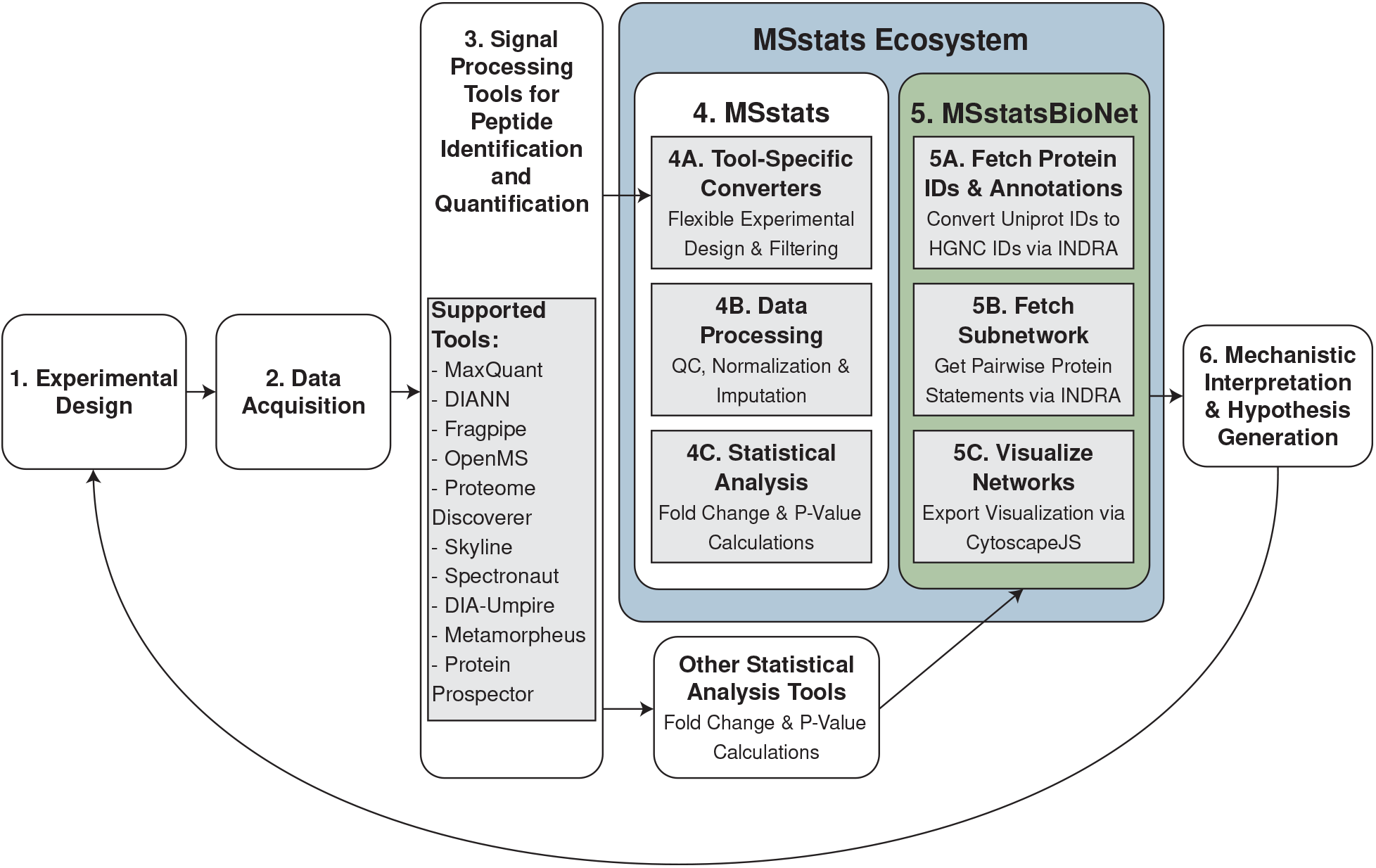
Summary of a typical data collection and analysis workflow with MSstats. MSstatsBioNet streamlines the step between ascertaining significance and formulating hypotheses for future experimental validation, by providing an R-based and graphical user interface to prior knowledge biological networks from INDRA.

After data acquisition (step 2 of **Figure 1**), the next step is to identify and quantify proteins in the acquired MS spectra (step 3 of **Figure 1**). A variety of spectral processing tools handle this task, including Spectronaut, ^56^ DIANN,^57^ OpenMS,^58^ MaxQuant,^59^ Metamorpheus,^60^ Fragpipe,^61^ Proteome Discoverer (Thermo Scientific), Skyline,^62^ DIA-Umpire,^63^ or Protein Prospector.^64^ These tools quantify protein “features” in the spectra. Here, we define a “feature” as a fragment ion for DIA, transition for SRM, and precursor for DDA.

### MSstats

MSstats offers an ecosystem of packages for differential abundance analysis of mass spectrometry-based proteomics experiments. MSstats offers functionalities that convert the outputs of data processing tools into a standardized format for data processing and statistical analysis (step 4A of **Figure 1**). Using MSstats, users can summarize feature-level intensities into summarized protein-level abundances while minimizing biases from technical artifacts via built-in methods for normalization, missing value imputation, and selection of high-quality features (step 4B of **Figure 1**). With summarized protein-level abundances, MSstats can perform differential analysis in a way that reflects the experimental design, leading to fold change estimates and p-value calculations (step 4C of **Figure 1**).

Here, we focus on the integration of statistical results with prior knowledge of biological networks (step 5 of **Figure 1**) to bridge statistical results with mechanistic interpretation and hypothesis generation for future experiments (step 6 of **Figure 1**). Once users formulate a hypothesis that may explain the statistical results, they can design a new experiment to validate the hypothesis, closing the experimental loop. This new experiment could involve a more targeted assay, such as an SRM assay to validate results from a DIA study, or an assay to measure a different data modality, such as a phosphoproteomics experiment to validate signaling activity inferred from results of a proteome-level DIA experiment.

### INDRA

INDRA is an automated software system that collects mechanistic biological relationships extracted from the biomedical literature by text mining systems together with relationships collected from manually curated databases. Each relationship is represented as an INDRA *statement*, a machine-readable representation of a mechanistic relationship between biological entities (e.g., “ATR activates RAD51”, “RPA1 binds to RPA2”, “MAPK3 phosphorylates ARRB1 at Serine 412”), characterized by a statement type (e.g., Phosphorylation, Complex, Activation, or Inhibition).^65^ INDRA refers to the biological entities that serve as the arguments of a statement as *agents*, which include gene products, protein families and complexes, small molecules, biological processes, and diseases. Each statement is supported by one or more pieces of evidence, where each piece of evidence is contributed by a single source and corresponds either to a single extraction from a sentence in a publication by a text mining system or to a single entry in a curated database. The text mining sources include Reach,^40^ Sparser,^41^ and RLIMS-P,^42^ and the curated database sources include BioGRID,^66^ SIGNOR,^67,68^ and Reactome.^36^

Because the same relationship is frequently reported by multiple sources and described in heterogeneous ways, INDRA performs a process of assembly. ^45^ Assembly first grounds each agent to standardized ontology identifiers with Gilda,^69^ so that different names referring to the same agent are normalized to a common reference. It then detects full and partial overlaps among statements and merges them into a single, non-redundant representation of each interaction or regulation, aggregating all of its supporting evidence across sources. As a result, each assembled statement carries an evidence count, the total number of supporting evidence instances across all sources, which serves as a proxy for how extensively studied a given relationship is in the literature. Beyond harmonizing individual statements, INDRA can also assemble a collection of statements into custom models and networks in a range of output formats, including executable rule-based models and various network representations.^65^

Running this assembly at scale over all accessible literature and a number of major pathway databases yields the INDRA Database (INDRA DB), a queryable resource of approximately 16 million assembled statements together with their aggregated evidence. A graph database-backed service for interacting with the INDRA DB statements is available at discovery.indra.bio, providing both a web service with an application programming interface (the INDRA API) and a graphical user interface (the INDRA web portal). The INDRA API supports programmatic path and network queries, including a subnetwork query, which retrieves all assembled statements among a set of input agents, as well as queries that normalize input identifiers (for example, mapping UniProt accessions and mnemonic identifiers to a standardized HGNC format) prior to a subnetwork query. The INDRA web portal links each statement to its supporting text-mined extractions and database entries, with sentences hyperlinked to the source publications, and allows users to curate individual pieces of evidence for correctness. Cloud-based automated reading and assembly workflows allow INDRA to be continuously updated, ensuring that the INDRA DB reflects the latest state of the literature.

Here, we integrate the subnetwork and identifier-normalization queries provided by the INDRA API with differential abundance analysis results from MSstats ecosystem packages, including MSstats, MSstatsTMT, and MSstatsPTM. This integration provides a streamlined interface to extract subnetworks among differentially abundant proteins/phosphosites and explore the associated literature for mechanistic interpretation and hypothesis generation for future follow-up experiments.

## Methods

### Overview

**Figure 2** shows the proposed workflow. MSstatsBioNet takes as input a list of differentially abundant proteins/phosphosites, their adjusted p-values (Benjamini-Hochberg FDR correction), and log_2_ fold changes produced by statistical analysis tools, specifically the MSstats ecosystem of packages such as MSstats,^12,13^ MSstatsTMT,^22^ and MSstatsPTM.^23^ MSstatsBioNet also provides an option for users to start with a table of proteins/phosphosites and their fold changes and p-values without orchestrating the full MSstats workflow.

**Figure 2:**
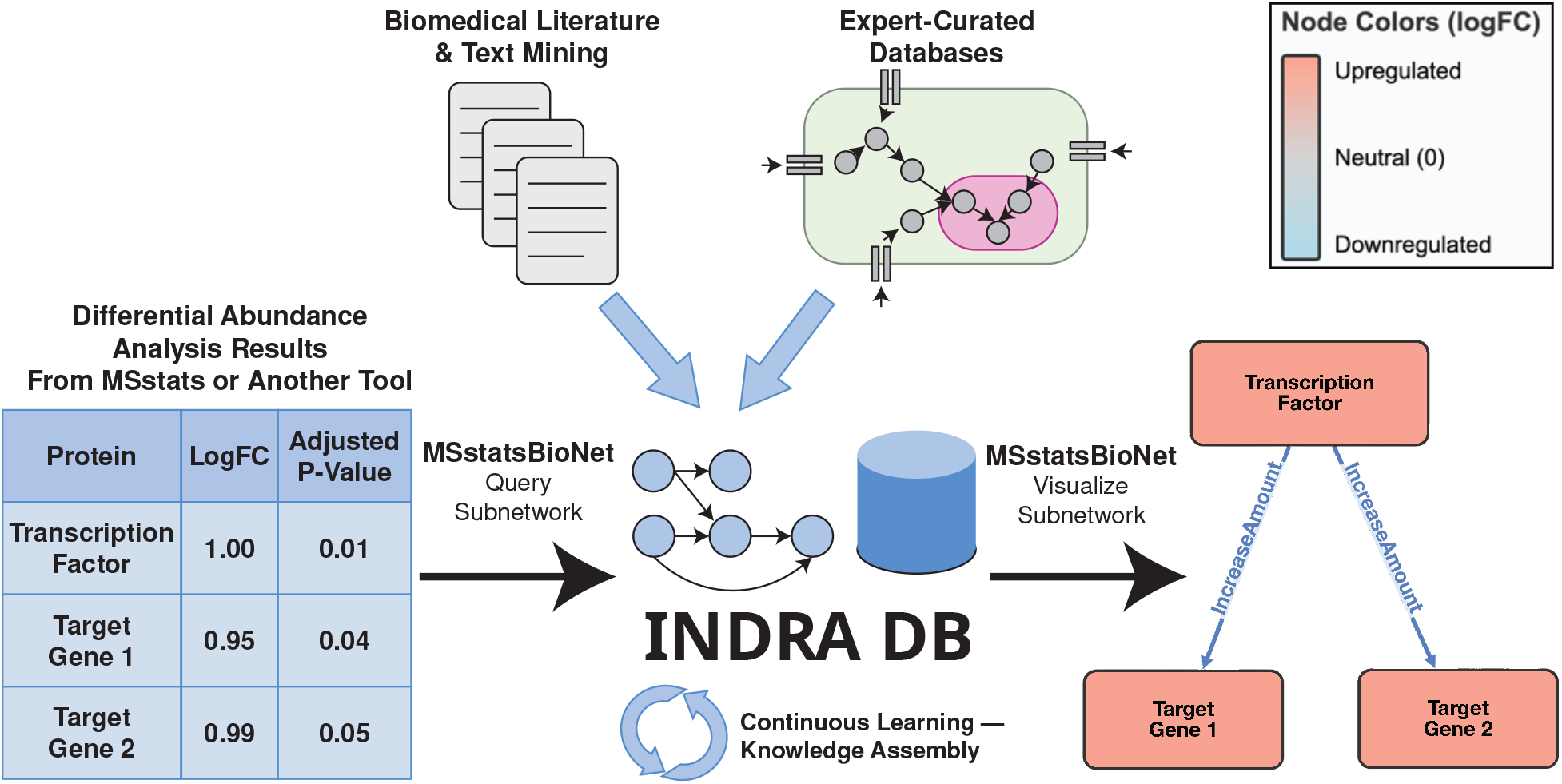
Querying workflow with MSstatsBioNet and INDRA. INDRA provides an interface for users to query a subnetwork found in curated databases and prior literature. After obtaining a set of mechanistic relationships, users can visualize the relationships as a network with MSstatsBioNet, where proteins are represented as nodes and mechanistic relationships are represented as edges. With the network, users can interrogate primary literature evidence to construct granular mechanistic narratives for iterative hypothesis generation.

Using a list of differentially abundant proteins, MSstatsBioNet outputs a subnetwork from INDRA and visualizes the results. A subnetwork is constructed as follows: for each pair of proteins in a list of proteins, INDRA checks if there exists a mechanistic relationship between the two proteins. If a relationship exists, INDRA returns that relationship to the user. In addition to fold change and p-value cutoffs for filtering, MSstatsBioNet offers additional filtering strategies summarized in **Supplementary Table S1**.

MSstatsBioNet provides a network visualization suite that allows users to dynamically explore the subnetwork and links users to the INDRA web portal. Here, users can inspect and curate the supporting evidence for each statement to construct a granular mechanistic narrative, empowering users to formulate a hypothesis for a plausible reason for the results.

### Fetch protein IDs and annotations

Mass spectrometry datasets vary in the types of identifiers used to represent proteins. To handle this heterogeneity, MSstatsBioNet queries the INDRA API to convert identifiers to HGNC IDs (step 5A of **Figure 1**), which is the required format for subsequent subnetwork queries. MSstatsBioNet accepts two common identifier formats as input: UniProt accession IDs and UniProt mnemonic IDs. PTM-containing identifiers are automatically parsed to extract the base protein identifier and the modification site, enabling site-level information to be carried through to downstream analyses. Along with ID conversion, MSstatsBioNet retrieves functional annotations for each protein from the INDRA API, indicating whether each protein is a kinase, phosphatase, or transcription factor, which allows users to filter subsequent subnetwork queries by protein functional class.

### Fetch subnetwork

MSstatsBioNet queries the INDRA API to retrieve a subnetwork of pairwise statements among the set of differentially abundant proteins or PTMs (step 5B of **Figure 1**). Prior to querying, users can restrict the input protein list by applying thresholds on adjusted p-value, absolute fold change, and direction of regulation (up-regulated only, down-regulated only, or both). Proteins detected in only one condition, which are represented as infinite log_2_ fold change values, can be optionally included under the assumption that their absence in the other condition reflects low abundance. Additional biological entities absent in the differential abundance results, such as chemical perturbations of interest or their known main targets, can be appended to the query by providing an identifier and namespace (e.g. HGNC:1925, CHEBI:28748).

After querying INDRA, the returned statements are further refined using several evidence-based filters. Users can set a minimum threshold on the evidence count, restricting the subnetwork to well-supported statements. Statements can also be filtered by statement type (e.g., Phosphorylation, Complex, Activation, or Inhibition) and by the source that contributed evidence. For PTM analyses, users can restrict statements to those where the specific modification site reported by INDRA matches a site that is also significant in the differential abundance results, reducing the search space when many PTMs are significant. Finally, users can incorporate manual curation performed on the INDRA web portal, where individual pieces of evidence flagged as incorrect are subtracted from each statement’s evidence count; statements that fall below the minimum evidence count threshold as a result are removed from the subnetwork. A full description of all available filtering options is provided in **Supplementary Table S1**.

MSstatsBioNet returns a table of retrieved statements, where each row represents a statement between two proteins along with metadata including the evidence count, statement type, source, and a link to the corresponding entry in the INDRA web portal. Users can follow the INDRA web portal link to inspect and interpret the underlying evidence in the context of their experiment. The portal includes the exact sentences that were used as evidence for a statement, along with a link to the full PubMed article. Because automated text mining can introduce errors, users can manually flag incorrect extractions on the INDRA web portal for a specific piece of evidence. After curation, the subnetwork query can be re-run with the curation filter enabled to automatically exclude the flagged statements.

### Visualize networks

MSstatsBioNet provides an interactive network visualization interface built on CytoscapeJS^70^ and htmlwidgets^71^ (step 5C of **Figure 1**). Cytoscape^72^ is a free, open-source software platform used for visualizing and analyzing networks that includes the desktop application Cytoscape Desktop and the Javascript-based library CytoscapeJS.^70^ htmlwidgets is a framework that allows javascript visualizations to be rendered as a self-contained HTML widget that embeds directly in R Markdown documents, RShiny applications, and standalone HTML files without requiring any external desktop software such as Cytoscape Desktop. Using both CytoscapeJS and htmlwidgets was a deliberate design choice: because the entire visualization runs in the browser via JavaScript, it is fully compatible with cloud-hosted deployments of RShiny, where connecting to a local desktop application is not possible.

Nodes are colored along a continuous blue–grey–red gradient according to the log_2_ fold change of the corresponding protein: down-regulated proteins are shown in blue, up-regulated proteins in red, and proteins included in the query but not measured in the experiment in grey. Edges are visually encoded by statement type: protein complex membership is shown as undirected lines while regulatory statements are shown as directed arrows colored by statement type. Where a protein has multiple significant PTM sites, each site is displayed as a small satellite node attached to the parent node. Hovering over an edge surfaces any overlap between the modification site reported by INDRA and the significant PTM sites in the input data. Each edge is hyperlinked to its corresponding INDRA web portal entry, enabling users to quickly inspect the supporting literature evidence. Right clicking an edge deletes the edge from the network visualization, which is useful for removing edges that do not align with the context of an experiment but are not necessarily text mining errors.

Networks can be exported as self-contained HTML files that bundle all visualization dependencies inline, allowing them to be shared with collaborators and opened in any web browser without additional software. The CytoscapeJS visualization enables MSstatsBioNet to be embedded into MSstatsShiny, which is a graphical user interface that allows users without programming expertise to access MSstats and its family of open-source packages and enables reproducibility through reanalysis scripts and visualization files for data sharing. ^54^ **Supplementary Figure S1** shows a screenshot of the new network interpretation tab in MSstatsShiny that integrates with MSstatsBioNet and **Supplementary Table S2** describes advantages and disadvantages of using MSstatsShiny over MSstatsBioNet R code.

## Experimental Datasets

The three datasets below demonstrate the ability of MSstatsBioNet to provide mechanistic interpretation of mass spectrometry-based differential protein and PTM abundance analysis results. The first case study investigated the effects of small molecule inhibitors on protein chromatin-binding in the THP-1 cell line. The second case study investigated the shared effects of doxorubicin in three different Rhabdomyosarcoma (RMS) cell lines. The third case study investigated the effects of Interferon-gamma (IFN*γ*) stimulation and *LGALS1* (galectin-1) knockdown in glomerular microvascular endothelial cells (GMECs),^73^ highlighting a potential link between clathrin-mediated endocytosis and IFN*γ* stimulation response. These case studies involve perturbations with some expected biological responses. However, MSstatsBioNet can also support comparisons that do not involve perturbations, such as disease vs. control analyses. **Table 1** provides a summary of the 3 datasets.

**Table 1:**
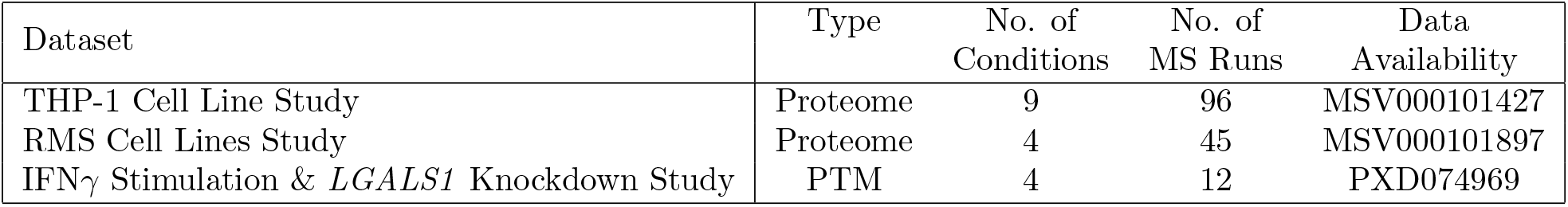
Type indicates whether the study measures global proteome or PTM abundance. No. of MS Runs indicates the number of mass spectrometry runs across all batches in the study. The data is available in MassIVE74 and PRIDE.^75^.

| Dataset | Type | No. of Conditions | No. of MS Runs | Data Availability |
| --- | --- | --- | --- | --- |
| THP-1 Cell Line Study | Proteome | 9 | 96 | MSV000101427 |
| RMS Cell Lines Study | Proteome | 4 | 45 | MSV000101897 |
| $\text{IFN}\gamma$ Stimulation & *LGALS1* Knockdown Study | PTM | 4 | 12 | PXD074969 |

### Dataset 1: THP-1 cell line study

#### Objective

In this case study, the objective was to assess the effect of small molecule compounds on the chromatin-binding activity of transcription factors and other nuclear proteins. In contrast to traditional chemotherapy drugs, small molecule inhibitors have emerged as an alternative treatment option for cancer and other diseases due to being more specific and targeted.^76^ Because transcription factors are heavily implicated in human cancer,^77^ understanding transcription factor and nuclear protein chromatin-binding behavior is important to uncover the therapeutic mechanisms of small molecules.

#### Experimental procedures and data processing

Intact, live cells from the THP-1 cell line were treated for four hours with one of eight small molecule compounds. A summary of the dosage, number of replicates, and known targets for each compound can be found in **Supplementary Table S3**. Then nuclei were isolated and the DNA-bound proteins were enriched. Chromatin-bound proteins were quantified using data independent acquisition (DIA) with Bruker timsTOF Ultra while identification and quantification were performed with the DIA-NN (version 1.8.1).^57^ The following search parameters for DIA-NN were changed from default: mass acc automatic = false; diann normalize = false.

#### Differential analysis

Differential analysis was performed through an MSstats pipeline. Since this was an exploratory study, the log intensities were normalized using median normalization. The data were summarized using top 100 features via Tukey’s Median Polish.^78^ The summarized protein abundances from all experimental conditions were fit into a single linear mixed-effects model. Differential analysis compared each small molecule treatment to DMSO.

### Dataset 2: Rhabdomyosarcoma (RMS) cell lines study

#### Objective

Topoisomerases (Topo) are crucial for maintaining proper cell division and proliferation by ensuring that DNA replication occurs smoothly.^79,80^ TopoI and TopoII are important targets in cancer therapy, as inhibiting these enzymes can lead to apoptosis in rapidly dividing cells like those in RMS. ^81–83^ While topoisomerase inhibitors are commonly used to treat various cancers, their toxic side effects are challenging to overcome.^84^

One approach to overcoming toxic side effects is to design analogues of these inhibitors. For example, while doxorubicin can drive apoptosis via both chromatin damage (histone eviction from open chromatin) and DNA damage (double-strand breaks induced by topo II poisoning),^85,86^ dimethyl-doxorubicin was designed to induce chromatin damage without invoking DNA damage,^86–89^ with the intention of reducing the cardiotoxic effects typically seen from doxorubicin’s DNA damage effects. Here, the objective was to disentangle the effects of chromatin damage, DNA damage, and their combination in RMS cell lines.

#### Experimental procedures and data processing

Intact, live cells from the RD, RH4, and RH30 cell lines were treated in triplicate for 2 hours at a 5*µ*M dose with dimethyl-doxorubicin, doxorubicin, or etoposide, or with DMSO control. Dimethyl-doxorubicin was used to isolate the effects of chromatin damage alone, etoposide was used to isolate the effects of DNA damage alone, and doxorubicin was used to assess their combined effects. RH4 and RH30 are human alveolar rhabdomyosarcoma cell lines derived from pediatric soft tissue tumors while RD is an embryonal rhabdomyosarcoma cell line. Nuclei of cells were isolated after 2 hours of drug exposure and the chromatin-bound proteins were enriched. Chromatin-bound proteins were quantified using data independent acquisition (DIA) with Bruker timsTOF SCP while identification and quantification were performed with DIA-NN. Mass spectrometry data files were analyzed by DIA-NN (version 1.8.1) using the quantms Nextflow pipeline.^90^ Library free search was performed using a human FASTA from Uniprot including isoforms (accessed 2022-03-15) and supplemented with common contaminants. The following search parameters for DIA-NN were changed from default: mass acc automatic = false; diann normalize = false.

#### Differential analysis

Since each cell line was processed as a separate batch rather than randomized across batches, a separate differential abundance analysis was performed for each cell line through an MSstats pipeline. Within each cell line, log intensities were normalized using median normalization and summarized using top 100 features via Tukey’s Median Polish.^78^ With summarized protein abundances, data were fit across all perturbation conditions and DMSO into a single linear mixed-effects model per cell line.

### Dataset 3: IFN*γ* stimulation and *LGALS1* knockdown study

#### Objective

Boshart et al. examined the effect of *LGALS1* knockdown on cell signaling in glomerular microvascular endothelial cells (GMECs) in the context of IFN*γ* stimulation.^73^ Antibody-mediated rejection (ABMR) accounts for *>*50 percent of premature kidney graft loss.^73^ ABMR is characterized by microvascular inflammation and injury to the kidney endothelium, which can be caused by IFN*γ*. In past studies, LGALS1 (galectin-1) was identified as significantly increased in ABMR and its expression was found to be decreased by IFN*γ* in GMECs in vitro.^39^ To investigate the effects of *LGALS1* knockdown, GMECs were exposed to one of four conditions: (1) *LGALS1* siRNA + IFN*γ*, (2) *LGALS1* siRNA + vehicle, (3) non-targeting control + IFN*γ*, and (4) non-targeting control + vehicle.

#### Experimental procedures and data processing

The experimental procedures and data processing were previously described in Boshart et al..^73^ Briefly, phosphoproteomics and proteomics data were acquired via data-dependent acquisition. Identification and quantification were performed with MaxQuant.

#### Differential analysis

Data were processed with MSstatsPTM with a global profiling adjustment. Log intensities were normalized using median normalization and summarized using top 100 features via Tukey’s Median Polish.^78^ With summarized protein abundances, data were fit across all experimental conditions into two linear mixed-effects models, one for the phosphoproteomics data and one for the global profiling proteomics data. Changes in phosphosite abundance were then adjusted by their corresponding changes in overall protein abundance as outlined in Kohler et al.. ^23^

## Results

### THP-1 cell line study

#### MSstatsBioNet extracted perturbation-specific subnetworks when differentially abundant proteins were detected

The comparison against DMSO produced 1 differentially abundant protein for K975, 2 differentially abundant proteins for PF477736, 3 differentially abundant proteins for DbET6, and 4 differentially abundant proteins for Jakafi with adjusted p-value ≤ 0.05. We observed no differentially abundant proteins among all other small molecules. For the initial subnetwork search, we used differential abundance results from MSstats to retrieve a subnetwork using MSstatsBioNet, filtering by adjusted p-value ≤ 0.05. If a small molecule had established targets, we included those in the subnetwork search, even if they were neither differentially abundant nor measured. Each statement was manually reviewed for technical extraction accuracy and relevance to our experimental context.

**Supplementary Figure S2** shows the initial uncurated subnetwork results on Jakafi, DbET6, K975, and PF477736. For positive control DbET6, we observed literature around how BRD2/3/4 may interact with each other. For other compounds, subnetwork search results varied. For K975, there were no edges between TEAD1-4 and the only differentially abundant protein TRMT6. For Jakafi, while we observed no literature evidence connecting the differentially abundant proteins to JAK1/2, we observed one protein complex edge between JAK2 and CUL4B from BioGRID,^66^ a database containing potential protein interactions from results of high-throughput experiments, such as MS affinity purification. For PF477736, we observed literature-backed evidence connecting the main target, CHEK1, to the differentially abundant proteins RPA1 and RPA2, as well as literature-backed evidence connecting RPA1 and RPA2. We chose to further investigate the literature behind the subnetwork produced when comparing PF477736 to DMSO since it was the only treatment where we observed literature-backed evidence connecting the main target to the differentially abundant proteins.

#### Initial subnetwork extraction revealed CHEK1-RPA phosphorylation relationships

When comparing PF477736 treated samples against DMSO, we observed 2 differentially abundant proteins, as shown in **Figure 3A**. **Figure 3B** shows the curated subnetwork found in INDRA among 2 differentially abundant proteins and CHEK1, filtered by Phosphorylation statements and INDRA evidence count ≥ 5 to ensure we initially reviewed literature relevant to our experimental context. We included CHEK1 in our subnetwork search because previous literature reported PF477736 may inhibit the phosphorylation activity of CHEK1.^91,92^ The subnetwork revealed that CHEK1 could phosphorylate both RPA1 and RPA2.

**Figure 3:**
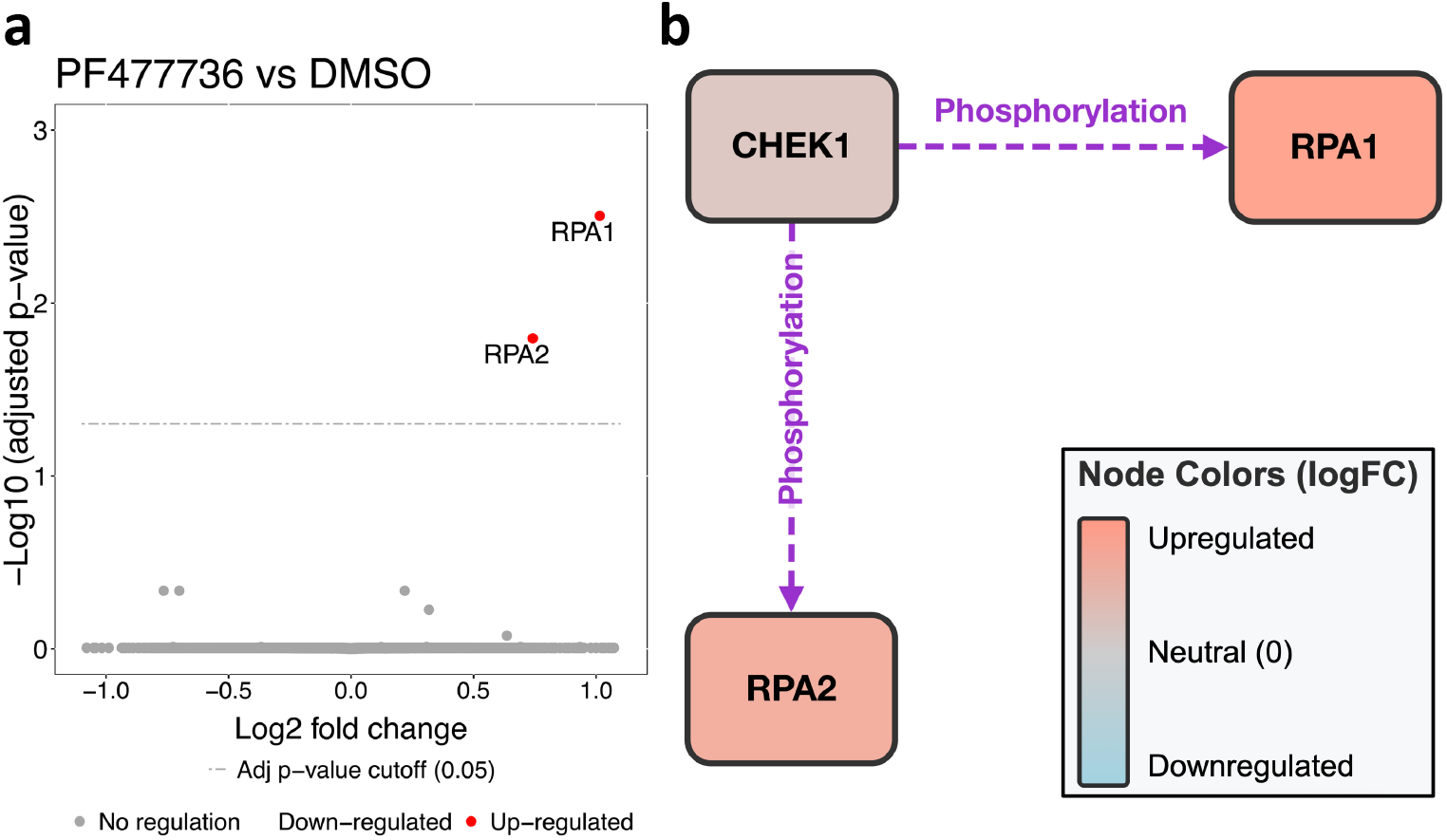
THP-1 Cell Line: Volcano plot and its subnetwork. a) Volcano plot with significant up-regulated changes in red. b) Subnetwork after manual curation with differentially abundant proteins and CHEK1, color coded by direction of regulation

As seen in **Supplementary Figure S3A**, we observed that according to Liu et al., CHEK1 phosphorylation of RPA1 may decrease RPA1 binding to single-strand DNA (ss-DNA) in the context of DNA damage repair.^93^ Taking the reverse of the reported effect, we inferred that inhibiting CHEK1 phosphorylation may then lead to more RPA1 binding to ssDNA. Considering that our experiment measured chromatin-bound proteins, this literature evidence was consistent with the up-regulation of RPA1 in the current study. Similarly, CHEK1-mediated phosphorylation of RPA2 may increase RPA2 binding to ssDNA,^94^ which was consistent with the up-regulation of RPA2 in the current study.

In a follow up study, we propose validating this finding via a phosphoproteomics experiment while inhibiting CHEK1 to demonstrate decreased phosphorylation of RPA1 and

RPA2.

#### Second subnetwork with RPA3 added additional support of a heterotrimeric DNA damage repair mechanism

DNA damage repair literature described that RPA1 and RPA2 could form a protein complex. This relationship was extracted by INDRA (**Supplementary Figure S4A**), which shows the text mined evidence for the statement “RPA1 binds with RPA2 and RPA3”, which is also counted as evidence for the statement “RPA1 binds RPA2”. This information suggests that RPA1, RPA2, and RPA3 may form a heterotrimeric complex.

**Supplementary Figure S4B** demonstrates the results of a subnetwork search with RPA3 and additionally included Protein Complex statements. We observed literature describing this heterotrimeric complex ^95–97^ and noted that RPA3 was also up-regulated in our results (p-value ≤ 0.05), although not differentially abundant at adjusted p-value ≤ 0.05. Although RPA3 was not significantly up-regulated, the lack of significance may have originated from a combination of an underpowered experiment and multiple hypothesis testing correction. Collectively, we hypothesize that inhibiting CHEK1 in THP-1 cells disrupted DNA damage repair checkpoints that involved removing RPA heterotrimeric complex off ssDNA via phosphorylation.

### Rhabdomyosarcoma (RMS) cell lines study

#### MSstatsBioNet extracted cell line-specific subnetworks for each perturbation

For the initial subnetwork search, we used differential abundance results from MSstats to retrieve a subnetwork using MSstatsBioNet, filtering by adjusted p-value *<* 0.1 and INDRA evidence count ≥ 2 for each cell line and perturbation combination. **Supplementary Table S4** documents the number of differentially abundant proteins and number of INDRA statements extracted with MSstatsBioNet for each cell line and perturbation combination. As many as 618 differentially abundant proteins were detected corresponding to 8684 INDRA statements.

#### Initial subnetwork extraction across cell lines revealed multiple up-regulated protein complexes

To decrease the initial search space, we focused on common mechanisms across cell lines. Here, common mechanisms refer to statements that were returned by MSstatsBioNet across all 3 cell lines and that showed consistent direction of regulation for a given perturbation. For example, if for one cell line, we obtained the statement WDR43 (up-regulated) and UTP18 (up-regulated) forming a protein complex, this statement was only kept if the same direction of change was observed with significance across all other cell lines. Doxorubicin treated samples against DMSO was chosen as the main focus of this case study (chosen for clarity of illustration; similar results were observed with dimethyl-doxorubicin).

**Figure 4A-C** show differential abundance results from MSstats for the RH4, RH30, and RD cell lines for doxorubicin vs DMSO respectively. **Figure 4D** shows the subnetwork results when comparing doxorubicin treated samples against DMSO samples, filtering by statements found across all three cell lines with consistent direction of regulation. We observed a subnetwork consisting of eight differentially abundant proteins, all of which were up-regulated across all cell lines. The subnetwork extraction suggested the eight up-regulated proteins may form protein complexes, something that was not readily evident from their differential abundance analysis results.

**Figure 4:**
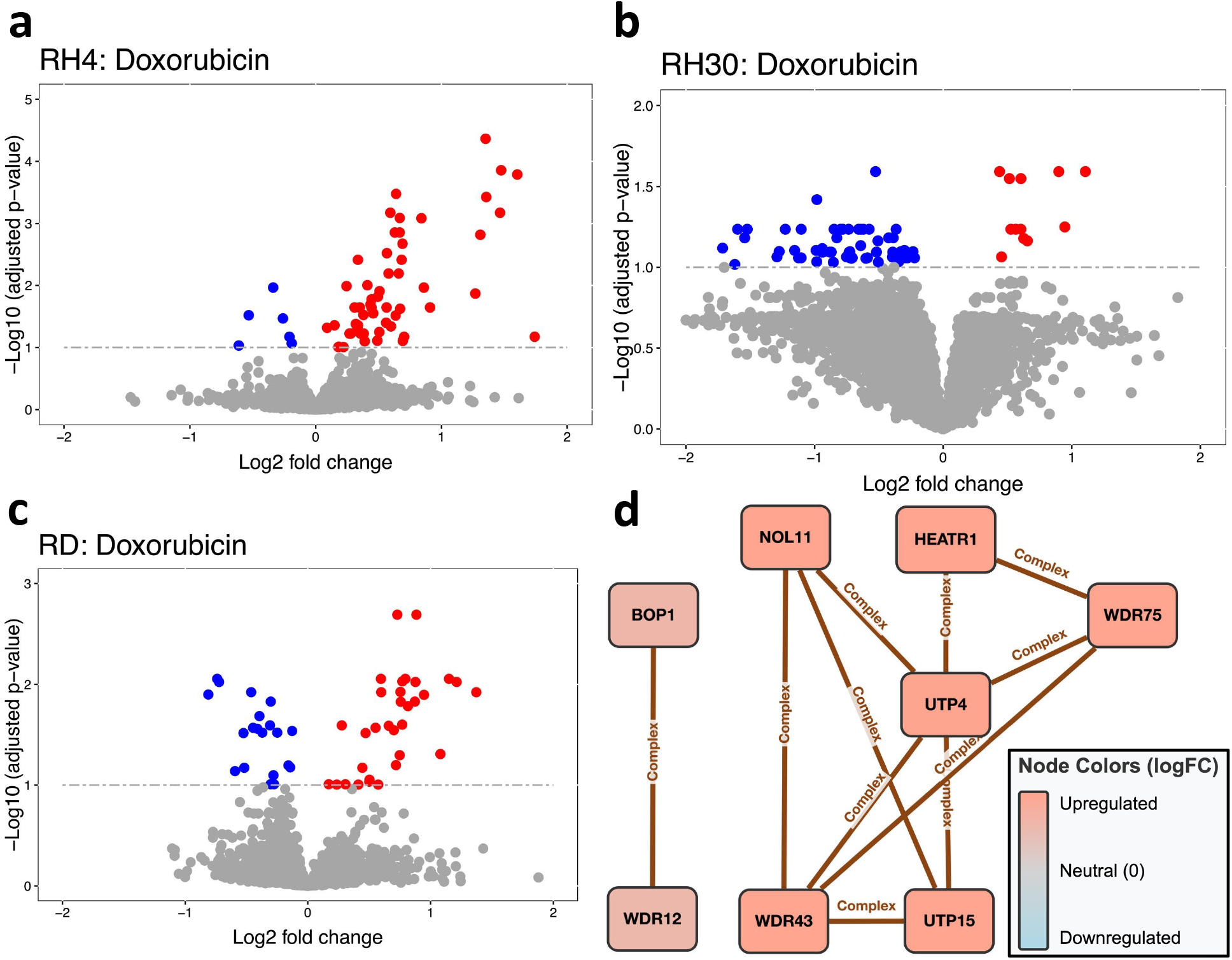
RMS Cell Lines Study: Volcano plots and corresponding subnetwork of common mechanisms across three cell lines. a) Volcano plot of doxorubicin compared to DMSO for the RH4 cell line, with differentially abundant proteins (adjusted p-value ≤ 0.1) in blue and red. b) Same as (a) except for the RH30 cell line. c) Same as (a) except for the RD cell line. d) Curated subnetwork containing statements that were returned by MSstatsBioNet across all 3 cell lines and that showed consistent direction of regulation. For example, the statement WDR43 and UTP18 was kept because we observed significant up-regulation of both proteins across all 3 cell lines. Proteins are color coded by direction of regulation and magnitude of median log2 fold change across cell lines..

#### Literature evidence characterized early ribosome assembly complexes UtpA and UtpB

We next sought to explain the up-regulation of the eight proteins in **Figure 4D**. We observed no mention of DNA damage or chromatin damage in the text-mined literature of the resultant subnetwork. Instead, we observed publications involving these protein complexes in ribosome biogenesis.^98–113^ We made similar observations of protein complexes involved in ribosome biogenesis when comparing dimethyl-doxorubicin with DMSO in **Supplementary Figure S5**. We noted that previous studies discussed the possibility that doxorubicin may affect ribosome biogenesis and induce nucleolar stress. ^114–116^

We first focused on explaining the regulation of the six-protein cluster consisting of UTP4, WDR43, WDR75, HEATR, UTP15, and NOL11 given this cluster consisted of more proteins and was up-regulated at a greater magnitude than the BOP1-WDR12 cluster. We noted that ribosome biogenesis occurs in the nucleolus, ^98^ and accounting for enrichment of chromatin-bound proteins, we considered different hypotheses depending on whether we assumed our assay captured only chromatin-bound proteins or also captured other nuclear artifacts, such as nucleolar proteins in close proximity.

If we assumed our assay only captured chromatin-bound proteins, then we considered whether translocation of these proteins from the nucleolus to the chromatin occurred. Fujimura et al. pointed out that heterotrimeric complex NOL11, UTP4, and WDR43 can be found in the nucleolus during interphase but can translocate to chromatin in the context of mitotic chromosome assembly. ^105^ While this context was different from the context of our study, where we expected mitosis to slow down due to chromatin damage from doxorubicin, this reference demonstrated that ribosome biogenesis factors could translocate from the nucleolus to chromatin upon a chromatin-related stimulus.

On the other hand, if we assumed our assay also captured other nuclear artifacts, such as nucleolar proteins, then we considered more granular hypotheses with respect to the stage of the ribosome biogenesis that was captured by the experiment. Most manuscripts in literature demonstrated that six of these proteins may be involved in early stages of ribosome biogenesis.^99,100,102–104,107^ Specifically, Hunziker et al. described a protein complex UtpA, consisting of UTP4, WDR43, WDR75, HEATR, UTP15, and UTP8 (a functional analog of NOL11), that could bind nascent pre-ribosomal RNA to initiate eukaryotic ribosome assembly. These six proteins in the UtpA protein complex matched the six-protein cluster in **Figure 4D**.

The literature also described a protein complex UtpB, consisting of PWP2, UTP18, WDR36, UTP6, WDR3, and TBL3, that may be similarly involved in binding of nascent pre-ribosomal RNA to initiate eukaryotic ribosome assembly.^102^ **Supplementary Figure S6** shows the expanded subnetwork when additionally including UtpB complex proteins. We observed that all UtpB complex proteins were up-regulated with adjusted p-value ≤ 0.3 across all cell lines, which matched the direction of regulation of the other UtpA complex proteins.

Furthermore, we observed in **Supplementary Figure S6** that proteins within the UtpA and UtpB complexes each showed internally consistent fold changes, yet the two complexes were clearly separated in their magnitudes of regulation, with the UtpA proteins showing greater magnitude. This result was relevant to insights from Hunziker et al., who demonstrated that UtpA complex may initiate ribosome assembly by binding to the nascent pre-rRNA, subsequently recruiting UtpB complex.^102^ As a result, we hypothesize that at the 2 hour timepoint of applying doxorubicin, we captured a snapshot of RMS cells in the middle of initiating ribosome assembly, specifically at the point when the cells were recruiting the UtpB protein complex to the nucleolus after already binding UtpA complex with nascent pre-rRNA. We reached the same conclusion when comparing dimethyl-doxorubicin to DMSO but not etoposide, the drug only making DNA double stranded breaks (data not shown).

#### Literature evidence described the PeBoW late-stage ribosome assembly complex

We observed multiple papers that described how BOP1, WDR12, and PES1 may form the PeBoW protein complex, which was documented as essential for assembly of large ribosomal subunit 60S.^108–113^ We initially hypothesized that BOP1 and WDR12 were up-regulated upon 2 hours of doxorubicin to facilitate later-stage maturation of the 60S ribosomal subunit. However, when adding PES1 to the subnetwork (not pictured), we observed no significant regulation of PES1 for any cell line. While we speculated the possibility for BOP1 and WDR12 to interact without PES1, PES1 has been documented as critical for the formation of the PeBoW complex and subsequent maturation of the large ribosomal subunit 60S.^108,110,111,113^ Building on our earlier observation that doxorubicin treatment may have captured RMS cells undergoing early ribosome assembly, we hypothesize that the absence of PES1 regulation may reflect a temporal gap: cells may have not yet progressed to the late-stage assembly step at which PES1 is recruited to form the PeBoW complex. This hypothesis could be tested by examining doxorubicin treatment at a later timepoint, at which PES1 regulation may become apparent if late-stage ribosome assembly is delayed rather than absent.

### IFN*γ* stimulation and *LGALS1* knockdown study

#### MSstatsBioNet extracted a protein-level subnetwork for multiple pairwise group comparisons

For the initial subnetwork search, we used differential abundance results from the adjusted model of MSstatsPTM to retrieve a subnetwork using MSstatsBioNet, filtering by adjusted p-value ≤ 0.2 and INDRA evidence count ≥ 1. **Supplementary Table S5** shows the differential abundance analysis results along with the number of INDRA statements extracted for each comparison. Across comparisons, we observed 1–10 differentially abundant phospho-sites. We observed 6 extracted INDRA statements when comparing non-targeting control + IFN*γ* vs non-targeting control + vehicle and 2 extracted INDRA statements when comparing *LGALS1* siRNA + vehicle vs non-targeting control + vehicle.

**Supplementary Figure S7** shows all INDRA statements extracted across all comparisons. We observed evidence that CAVIN1 and AHNAK could be part of the same protein complex.^117^ We also observed that *β*-arrestin-1 (ARRB1) may be in close proximity to PIK3C2A and SRRM2 based on high throughput MS affinity capture experiments from BioGRID.^66^ In both cases, the evidence provided no insight on whether these interactions could be triggered by a specific PTM event.

#### MSstatsBioNet extracted a site-specific subnetwork for each group comparison

To produce a site-specific subnetwork, we additionally filtered for phosphorylation statements where the phosphorylation site significantly regulated in the experiment matched the phosphorylation site described in INDRA. To increase our subnetwork search space knowing that there was limited literature studying site-specific interactions, we also included phosphosites in our subnetwork search that were missing in only one condition, since these were the most potentially important, signifying on-off signaling. Two such phosphosites of interest were mitogen-activated protein kinase 3 (MAPK3) phosphorylations at Threonine 202 and Tyrosine 204. These two phosphopeptides were absent in cells under non-targeting control + IFN*γ*. We assumed that any missing phosphosites were absent due to low abundance in that condition. We justified this assumption knowing that the data were acquired via data dependent acquisition, which selects ions for fragmentation based on MS1 abundance rank,^118^ meaning lower-abundance peptides are less likely to be sampled. To improve the interpretability of the extracted subnetworks, we also performed separate subnetwork searches for up- and down-regulated phosphosites, restricting each search to phosphosites with the same direction of regulation.

**Supplementary Table S6.** shows the number of missing phosphosites in one condition along with the number of site-specific INDRA statements extracted for each comparison. MSstatsBioNet extracted between 4–20 site-specific INDRA statements for each comparison. **Supplementary Figure S8A** shows an example of the site-specific subnetwork extracted when comparing non-targeting control + IFN*γ* vs non-targeting control + vehicle with these additional filters, restricted to down-regulated phosphosites.

#### Site-specific subnetwork revealed MAPK3-ARRB1 phosphorylation linked to clathrin-mediated endocytosis

We observed the most INDRA statements for the following two comparisons: Non-targeting control + IFN*γ* vs Non-targeting control + vehicle (examining the effect of IFN*γ* stimulation) and *LGALS1* siRNA + IFN*γ* vs Non-targeting control + IFN*γ* (examining the effect of LGALS1 knockdown). Within the subnetwork shown in **Supplementary Figure S8**, we chose to initially explore the potential phosphorylation relationship MAPK3 to ARRB1 since ARRB1 was the only protein in the subnetwork with phosphosites that were measured across all experimental conditions.

As shown in **Figure 5C**, under IFN*γ* stimulation, there was down-regulation of phosphorylated MAPK3 at Y204 and T202 and phosphorylated ARRB1 at S412 in the data. In the case of MAPK3, its phosphosites were not measured under IFN*γ* stimulation in the absence of *LGALS1* knockdown. When *LGALS1* was silenced followed by IFN*γ* stimulation, the direction of regulation was in the opposite direction as shown in **Figure 5D**, i.e. up-regulation of phosphorylated MAPK3 at Y204 and T202 and phosphorylated ARRB1 at S412. The direction of regulation of the phosphosites of MAPK3 was inferred based on an assumption that missingness was due to low abundance.

**Figure 5:**
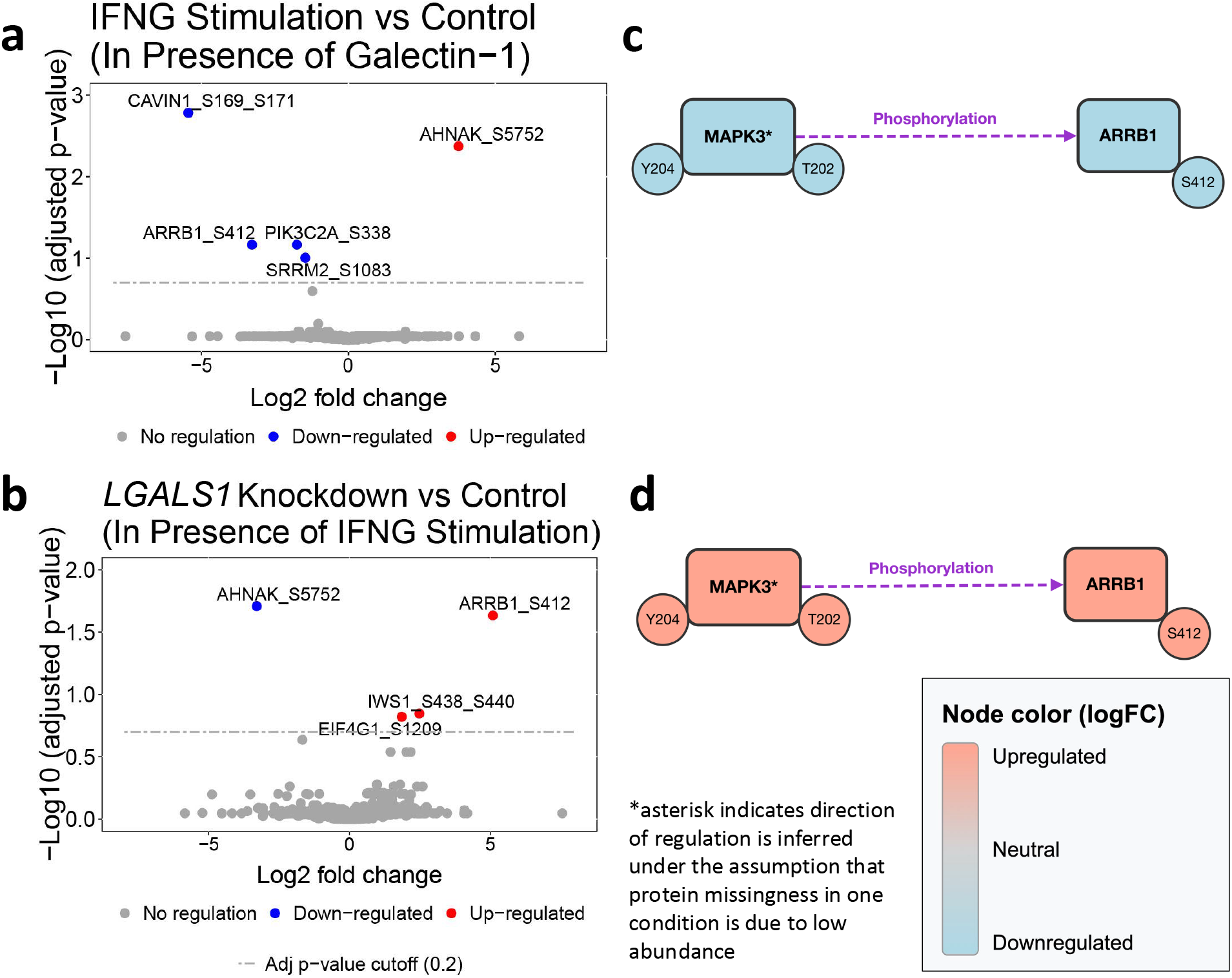
IFN*γ* Signaling Study: Volcano plot and subnetwork with significant phosphosites after manual curation. a) Volcano plot with significant phosphosites at adjusted p-value cutoff of 0.2 comparing GMECs under Non-targeting control + IFN*γ* stimulation vs Nontargeting control + vehicle (examining the effect of IFN*γ* stimulation). b) Volcano plot with significant phosphosites at 0.2 adjusted p-value cutoff comparing GMECs *LGALS1* siRNA + IFN*γ* vs Non-targeting control + IFN*γ* (examining the effect of LGALS1 knockdown). c) Subnetwork after manual curation comparing GMECs under Non-targeting control + IFN*γ* vs Non-targeting control + vehicle (examining the effect of IFN*γ* stimulation). Proteins are color coded by direction of regulation. MAPK3 at Threonine 202 and Tyrosine 204 is considered down-regulated since it is not measured upon IFN*γ* stimulation. d) Subnetwork after manual curation comparing GMECs under *LGALS1* siRNA + IFN*γ* vs Non-targeting control + IFN*γ* (examining the effect of LGALS1 knockdown). Proteins are color coded by direction of regulation. MAPK3 at Threonine 202 and Tyrosine 204 is considered up-regulated since it is not measured in Control.

**Supplementary Figure S8B** shows the site-specific text mined evidence extracted by INDRA for the MAPK3-ARRB1 phosphorylation statement. Looking through INDRA, we observed that all descriptions of the MAPK3-ARRB1 phosphorylation statement pointed to work done by Lin et al.,^119^ who used multiple assays to confirm that MAPK3 phosphorylates ARRB1 at S412. An in-vitro kinase assay was leveraged, where GST-MAPK3 with ATP were co-incubated leading to increased phosphorylation of wild type vs ARRB1 S412D mutant. In turn, MEK1 inhibition resulted in decreased ARRB1 S412 phosphorylation.^119^ Lin et al. characterized this phosphorylation behavior as part of a feedback loop where an agonist could lead to initial dephosphorylation of ARRB1 at serine 412, allowing ARRB1 to bind to SRC, triggering a signaling cascade involving *β*_2_-adrenergic receptor and clathrin-mediated endocytosis, leading to MAPK3 activation. As a negative feedback mechanism, Lin et al. suggested that MAPK3 phosphorylates ARRB1 at serine 412 to dissociate from SRC. Additional experiments between ARRB1 and SRC supported that ARRB1 and SRC binding may be required for MAPK3 activation via *β*_2_-adrenergic receptor internalization.^120,121^ Although SRC had no measured phosphopeptides, we confirmed it was measured in the global proteome dataset.

We determined this phosphorylation relationship as potentially relevant to this study because earlier studies linked clathrin-mediated endocytosis to signaling.^122–125^ Specific to IFN*γ* signaling, one study demonstrated that IFN*γ* receptors could be internalized via clathrin-mediated endocytic pathways.^122^ We additionally acknowledged that with the same dataset, Boshart et al. observed down-regulation of CAVIN1 at sites S169 and S171 upon IFN*γ* stimulation and *LGALS1* knockdown, and linked this down-regulation to a loss of co-localization with caveolin-1 (CAV1), an interaction involved in the formation of caveolae, which has been documented to mediate clathrin-independent endocytosis.^73^

Therefore, we propose that phosphorylation of ARRB1 at serine 412 was decreased in order to interact with SRC and trigger clathrin-mediated endocytosis. Observing that ARRB1 phosphorylation at serine 412 as well as MAPK3 phosphorylation at Y204 and T202 were down-regulated as a result of IFN*γ* stimulation, we suggest that IFN*γ* stimulation in GMECs indirectly triggered clathrin-mediated endocytosis. This suggests that MAPK3 was initially inactivated to prevent phosphorylation of ARRB1. Observing that *LGALS1* knockdown in presence of IFN*γ* stimulation led to an up-regulation of ARRB1 phosphorylation at S412 and MAPK3 phosphorylation, we suggest that galectin-1 was critical for ensuring clathrin-mediated endocytosis under IFN*γ* stimulation.

### Including IFN*γ* canonical pathway proteins in subnetwork search implicated STAT3 as a potential negative regulator of MAPK3

In **Supplementary Figure S9**, we produced a subnetwork that included proteins relevant to IFN*γ* stimulation to our subnetwork query, notably IFN*γ*, galectin-1, STAT1, and STAT3.^126^ We observed that in this subnetwork there existed an inhibition relationship from STAT3 to MAPK3. While MAPK3-specific evidence for this relationship was sparse, querying INDRA for statements at the level of the ERK family (ERK1/MAPK3 and ERK2/MAPK1) surfaced kidney-specific evidence that STAT3 inhibits ERK. In particular, INDRA provided evidence that, in mouse proximal tubular cells, STAT3 attenuates EGFR-mediated ERK activation during oxidant stress.^127^ We consequently inferred that STAT3 activation led to decreased MAPK3 phosphorylation.

Although we did not measure STAT3 phosphosites with mass spectrometry, we assumed based on prior literature that IFN*γ* stimulation led to increased STAT3 phosphorylation in this study. In addition, Boshart et al. previously performed a Western blotting of Phospho-STAT1 at tyrosine 701 after applying the same dose and duration of IFN*γ* treatment on GMECs and confirmed Phospho-STAT1 up-regulation,^73^ which suggested Phospho-STAT3 may also be up-regulated. Under this assumption, we hypothesize in **Supplementary Figure S9** that IFN*γ* stimulation resulted in increased STAT3 phosphorylation, decreasing MAPK3 phosphorylation, further attenuating ARRB1 phosphorylation. Reduced ARRB1 phosphorylation may have resulted in binding to SRC, triggering clathrin-mediated endocytosis.

Given INDRA also contained evidence that galectin-1 led to the phosphorylation of STAT3,^128,129^ we inferred that *LGALS1* knockdown decreased STAT3 phosphorylation, which led to higher phosphorylation of MAPK3, and more phosphorylation of ARRB1. Increased ARRB1 phosphorylation may have reduced binding to SRC and lowered clathrin-mediated endocytosis.

We also observed that INDRA contained a phosphorylation relationship from IFN*γ* to MAPK3 (not shown), suggesting that IFN*γ* stimulation could increase MAPK3 phosphorylation, which contrasted with the results of this study. When reading the text-mined literature of the IFN*γ* to MAPK3 relationship, we observed that the studies applied different doses of IFN*γ* and examined different cell lines,^130–132^ which both may be responsible for this difference.

## Discussion

MSstatsBioNet provides a streamlined interface to build a granular mechanistic narrative from context-specific prior knowledge networks, streamlining hypothesis generation for future experiments. Through three case studies, we demonstrated the ability of MSstatsBioNet to facilitate hypothesis generation. The process begins with a set of differentially abundant proteins supported by a prior knowledge subnetwork as illustrated in **Figure 2**, followed by iterative expansion of the subnetwork based on literature exploration.

In some cases, one may expect a relatively low number of differentially abundant proteins. For example, with the THP-1 cell line study, the limited number of differentially abundant proteins may not be surprising given the targeted nature of small molecule inhibitors. Additionally, 4 hours may be too short a time frame to observe transcriptional effects.^133^ Furthermore, since the experiment specifically measured chromatin-bound proteins, it did not necessarily reflect the overall protein abundance in a cell. Nevertheless, we were able to extract a context specific subnetwork that provided a mechanistic interpretation of the differential analysis results.

The proposed workflow has several limitations. First, understudied proteins can have a sparse prior literature.^134^ In these cases, one must rely on lower evidence statements which, for technical reasons, may have a higher chance of being incorrect. Similarly, annotations on PTM site-level interactions are sparse,^135^ and interpretable hypotheses often require multiple compounding assumptions. For example, with the IFN*γ* stimulation and *LGALS1* knockdown case study, we assumed that all the phosphosites of MAPK3 in one condition were missing for reasons of low abundance. But the phosphosites of MAPK3 could also be missing for other technical reasons, such as errors in the enrichment step, batch effects, or spectral misidentification. Second, MSstatsBioNet can only handle human proteins. While users could map proteins of other species to human analogues, this approach has limitations given that not every protein in another species maps to a human analogue.

Lastly, any text mining system will suffer from extraction inaccuracies. Common systematic extraction errors include mistaking a protein for a different biological entity, mistaking the directionality of a relationship between two entities, and, in the case of PTMs, specifying a wrong PTM site involved in a relationship.^45^ While MSstatsBioNet streamlines inspection and curation of evidence, inspecting every statement in a subnetwork becomes intractable when an experiment has many differentially abundant proteins, leading to a large network size to the point of being uninterpretable. While one could increase the INDRA evidence count threshold to reduce the size of the subnetwork, one risks losing access to novel hypotheses given the overstudied nature of highly cited statements. In the RMS cell lines study, we demonstrated one can search for common biological mechanisms across multiple similar conditions to narrow down the initial search space. Another approach to handle many differentially abundant proteins could involve performing enrichment analysis to provide high level terms and then constraining the subnetwork to sources with specific mentions of the terms. In this way, enrichment analysis can serve as a complementary tool for in-depth literature exploration with INDRA. The extracted subnetwork can also be refined using advanced downstream modeling in the spirit of NetworkCommons,^136^ PHONEMeS,^137^ COSMOS,^138^ and CORNETO,^139^ and we hope that this work will become a valuable resource for this type of modeling in the future.

Despite the limitations, MSstatsBioNet is applicable to proteomics and PTM experiments and broadly applicable for a wide variety of experiments. The open-source implementation makes the tool available to the proteomics community, allowing researchers to conduct reproducible data interpretation, share hypotheses, and build plausible and novel hypotheses for future experiments.

## Acknowledgement

The authors thank Talus for their contribution in testing the MSstatsBioNet package, Sarah Szvetecz for suggestions to this manuscript, and Sofia Farkona for processing the IFN*γ* signaling case study with MSstatsPTM. BMG and KK were supported by the DARPA ASKEM and ARPA-H BDF programs (HR00112220036).

## Supporting Information Available

Experimental data for all case studies can be found on MassIVE and PRIDE. Reanalysis code for the case studies are available in the corresponding MassIVE container of those datasets and on Zenodo.

