## Supplementary Figures and Tables for "MSstatsBioNet: Integrating Statistical Analyses with Prior Knowledge Biomolecular Networks for Quantitative Proteomics and Phosphoproteomics"

#### Supplementary Information

July 9, 2026

Anthony Wu[1], Devon Kohler[1], Pruthvi Prakash Navada[1], Julia E. Robbins[2], Gabriel E. Boyle[3], Alex Boshart[4,5], Klas Karis[1], Jacques Neefjes[8], Ana Konvalinka[4,5,6,7], Jay Sarthy[3], Lindsay Pino[2], Benjamin M. Gyori[1*], Olga Vitek[1*]

[1] Khoury College of Computer Science, Northeastern University, Boston, MA, USA 02115
[2] Talus Biosciences, Seattle, WA, USA 98122
[3] Seattle Children's Hospital, Seattle, WA, USA 98105
[4] Institute of Medical Science, University of Toronto, Toronto, ON, M5S 3H2, Canada
[5] Ajmera Transplant Centre, University Health Network, Toronto, ON, CA M5G 1L7, Canada
[6] Department of Medicine, Division of Nephrology, University Health Network, Toronto, ON, M5G 2N2, Canada
[7] Laboratory Medicine and Pathobiology, University of Toronto, Toronto, ON, M5S 3H2 Canada
[8] Department of Cell and Chemical Biology and Oncode Institute, Leiden University Medical Center, Leiden 2300 RC, The Netherlands

### S1 Methods

#### S1 Summary of the functionalities of MSstatsBioNet

Table S1: Summary of MSstatsBioNet functionalities

| Functionality | Description | Examples |
| --- | --- | --- |
| BH-Adjusted P-Value Filter | Only include proteins / PTMs that are significant under a specified significance threshold | Setting to 0.05 limits subnetwork to proteins / PTMs that are significant with adjusted p-value $\leq$ 0.05 |
| Log Fold Change Filter | Only include proteins / PTMs whose absolute log fold change is above a certain threshold | Setting to 1 limits subnetwork to proteins / PTMs with a log fold change $\geq$ 1 or log fold change $\leq$ -1 |
| Direction of Regulation | Only include proteins in subnetwork search that share the same direction of regulation. Useful for performing a subnetwork search of one side of a volcano plot | Setting to `"up"` narrows subnetwork search to only up-regulated proteins |
| Include Infinite Fold Change Proteins | Setting to TRUE includes proteins or PTMs with log fold change set as Infinity (missing in control) or -Infinity (missing in treatment). Assumes missingness is due to low abundance. | In a comparison of tumor vs. normal tissue, a protein detected only in tumor samples will have $\log_2 \mathrm{FC} = +\infty$. Setting to TRUE includes such proteins in the subnetwork search. |
| Evidence Count Filter | Only include edges supported by at least a specified number of evidence instances across all curated databases and text mining sources. When the curation filter is also enabled, manually flagged incorrect evidence is subtracted before this threshold is applied | Setting to 5 limits the subnetwork to edges with at least 5 supporting evidence instances |
| Statement Types Filter | Only include certain types of relationships | Setting to `c("Phosphorylation", "Complex")` limits edges to those describing phosphorylation events or protein complex / binding events |
| Sources Filter | Only include edges supported by certain text mining systems or curated databases | Setting to `c("reach", "phosphosite_plus")` limits edges to those found by the Reach text mining system and the PhosphoSitePlus curated database |
| Filter by PTM Site | Setting to TRUE only includes edges where the modification site reported by INDRA matches a PTM site that is also significant in the differential abundance results. Useful when there are many significant PTMs to reduce the initial exploration search space. | If MAPK3 is significant at any site and ARRB1 at S412 is significant in the input and INDRA reports an edge describing MAPK3 phosphorylation of ARRB1 at S412, that edge is retained. An edge describing phosphorylation of ARRB1 at a different site (or not specific to a site) would be removed. |
| Filter by Curation | Exclude edges for which manual curation on the INDRA web portal has flagged evidence as incorrect, where doing so reduces the net evidence count below the specified minimum | If an edge has 6 supporting evidence instances but 2 have been flagged as incorrect on INDRA DB, the net evidence count becomes 4. With an evidence count threshold of 5, this edge would be removed. |
| Additional Entities of Interest | A list of biological entities of interest to include in the search, whether or not they were measured. Entities are specified as namespace-prefixed identifiers | Setting to `"HGNC:1925"` ensures that CHEK1 is included in the subnetwork search. Setting to `"CHEBI:4911"` includes the chemical entity, Etoposide. |

### S2 MSstatsShiny integration with MSstatsBioNet

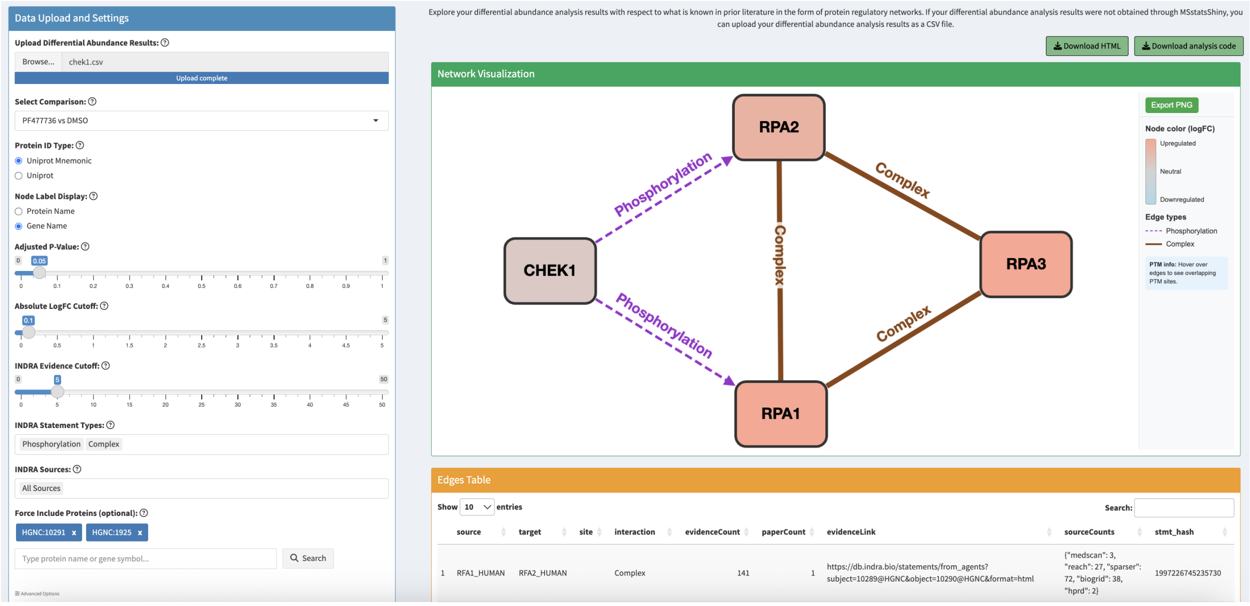

Figure S1: **THP-1 case study: Screenshot of MSstatsShiny app with new network interpretation tab.** The side panel consists of various MSstatsBioNet features, notably the BH-adjusted p-value filter, log fold change filter, evidence count filter, statement type filter, source filter, and additional proteins of interest to include in the subnetwork query. On the `Network Visualization` section, users can export the network as a PNG, download the network as an HTML, and download the MSstatsBioNet code to reproduce the network. The `Edges Table` consists of the edge metadata returned from MSstatsBioNet, which includes each relationship's statement type (interaction), evidence count (evidenceCount), link to INDRA (evidenceLink), and text mining systems / curated databases that extracted the relationship (sourceCounts).

Table S2: MSstatsBioNet R Code vs MSstatsShiny Integration

| Consideration | MSstatsBioNet R Code | MSstatsShiny Integration |
|---|---|---|
| Ease of Use | Not user friendly for users with limited programming experience | User friendly especially for those with limited programming experience |
| Multiple Dataset Integration | Custom code in conjunction with MSstatsBioNet R code can enable analysis and interpretation of multiple datasets | Only enables interpretation of one comparison of one dataset at a time |
| Manual Edge Addition and Deletion | Edges can be added and deleted at scale via R code | Edges can only be filtered out one at a time through manual curation of incorrect evidence on the INDRA web portal. |
| Variety of Filters | Includes the full set of MSstatsBioNet filters and metadata for subnetwork querying | Filters are limited to the most generalizable set of filters |

### S2 Results

#### S1 THP-1 cell line study: supplementary materials

Table S3: THP-1 cell line summary: a total of 96 samples were processed on one plate.

| Treatment | Sample Type | Dosage | Target | Replicates |
|---|---|---|---|---|
| DMSO | Negative control | n/a | n/a | 48 |
| K784-3670 | Treatment | 20$\mu$M | Unknown | 6 |
| VTP50469 | Treatment | 20$\mu$M | MEN1 | 6 |
| PF477736 | Treatment | 20$\mu$M | Chk1 | 6 |
| Jakafi | Treatment | 20$\mu$M | JAK2 | 6 |
| K975 | Treatment | 20$\mu$M | TEAD1 | 6 |
| VE-821 | Treatment | 20$\mu$M | ATR | 6 |
| K784-3183 | Treatment | 20$\mu$M | Unknown | 6 |
| dBET6 | Positive control | 50$\mu$M | BRD2/3/4 | 6 |

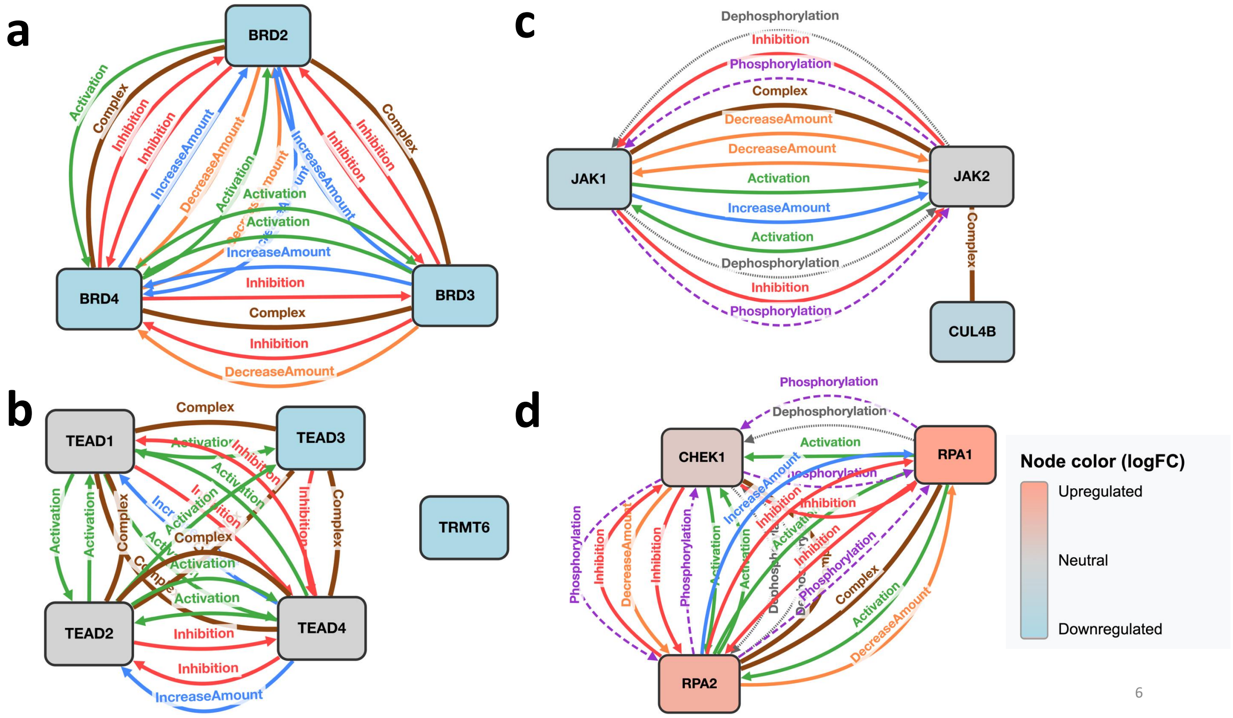

Figure S2: **Uncurated subnetwork extracted from INDRA for DbET6, K975, Jakafi, and PF477736. All grey-colored proteins were not measured.** a) Subnetwork search result of differentially abundant proteins of DbET6 vs DMSO. b) Subnetwork search result of single differentially abundant protein (TRMT6) of K975 vs DMSO comparison along with K975 main targets TEAD1-4. There exists no statement in INDRA between TRMT6 and TEAD1-4. c) Subnetwork search result of differentially abundant proteins of Jakafi vs DMSO along with Jakafi main targets JAK1-2. Protein complex statement between JAK2 and CUL4B is from a high-throughput affinity-capture MS experiment reported in BioGRID. d) Subnetwork search result of differentially abundant proteins of PF477736 vs DMSO along with the main target CHEK1.

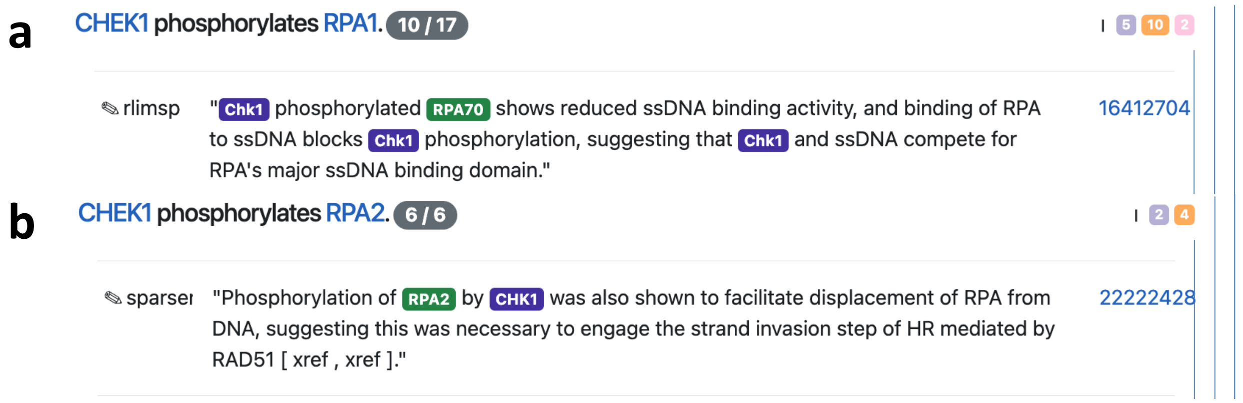

Figure S3: **Screenshot of INDRA web portal displaying extracted text mined evidence.** a) Screenshot of INDRA web portal with example text mined evidence for "CHEK1 phosphorylates RPA1" statement. CHK1 corresponds to CHEK1 and RPA70 is a synonym for RPA1. There is an evidence count of 17 for this "CHEK1 phosphorylates RPA1" relationship, notably 5 extractions from Reach, 10 extractions from Sparser, and 2 extractions from RLIMS-P. b) Screenshot of INDRA web portal with example text mined evidence for "CHEK1 phosphorylates RPA2" statement. c) Screenshot of INDRA web portal with example text mined evidence for "RPA2 phosphorylates CHEK1" statement.

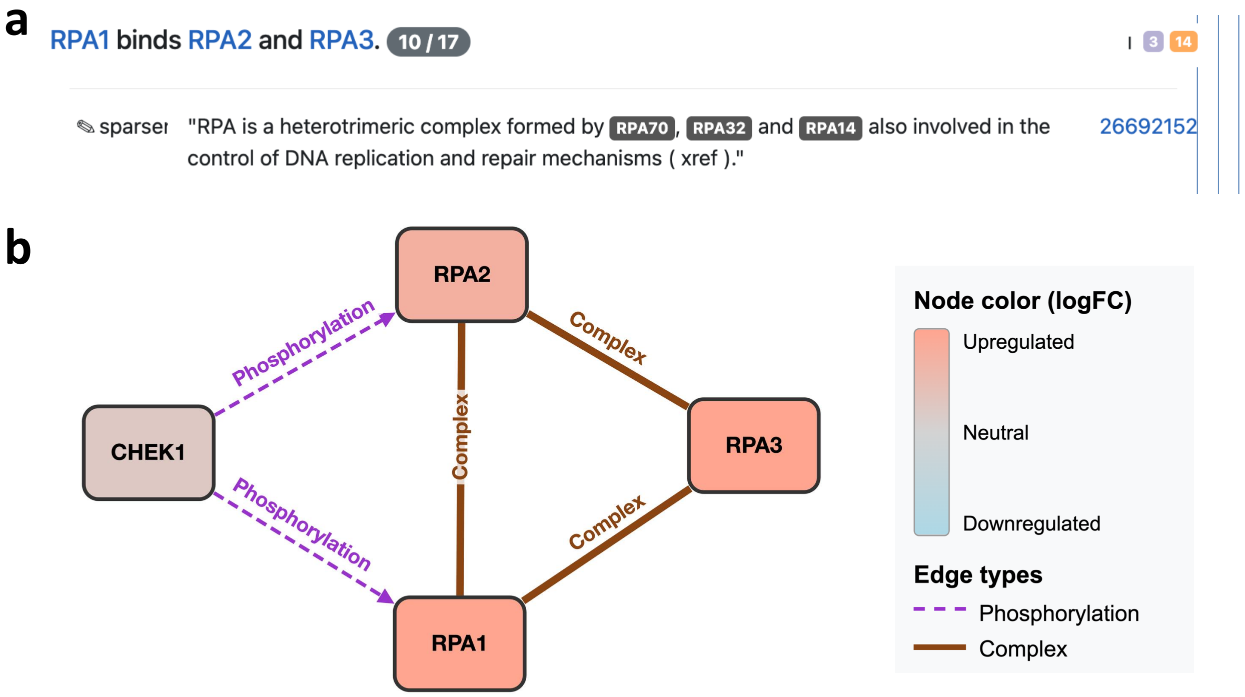

Figure S4: **Screenshot of INDRA web portal displaying extracted text mined evidence for RPA1 and RPA2 and corresponding subnetwork including RPA3.** a) Screenshot of INDRA web portal with example text mined evidence for "RPA1 binds RPA2 and RPA3" statement. b) Corresponding subnetwork with RPA3 forced to be included in the subnetwork.

#### S2 RMS cell lines study: supplementary materials

Table S4: RMS cell lines subnetwork extraction results (vs DMSO). No. of INDRA statements for dimethyl doxorubicin treatment on RH4 cells was calculated based on the 400 proteins with the lowest adjusted p-value since the INDRA subnetwork API is limited to querying 400 proteins at a time, representing a lower threshold on the number of INDRA statements.

| Treatment | Cell Line | No. Differentially Abundant Proteins | No. of INDRA Statements |
|---|---|---|---|
| Doxorubicin | RH4 | 61 | 487 |
| Doxorubicin | RH30 | 74 | 141 |
| Doxorubicin | RD | 52 | 234 |
| Dimethyl Doxorubicin | RH4 | 618 | 8684+ |
| Dimethyl Doxorubicin | RH30 | 26 | 83 |
| Dimethyl Doxorubicin | RD | 112 | 779 |
| Etoposide | RH4 | 12 | 64 |
| Etoposide | RH30 | 0 | 0 |
| Etoposide | RD | 0 | 0 |

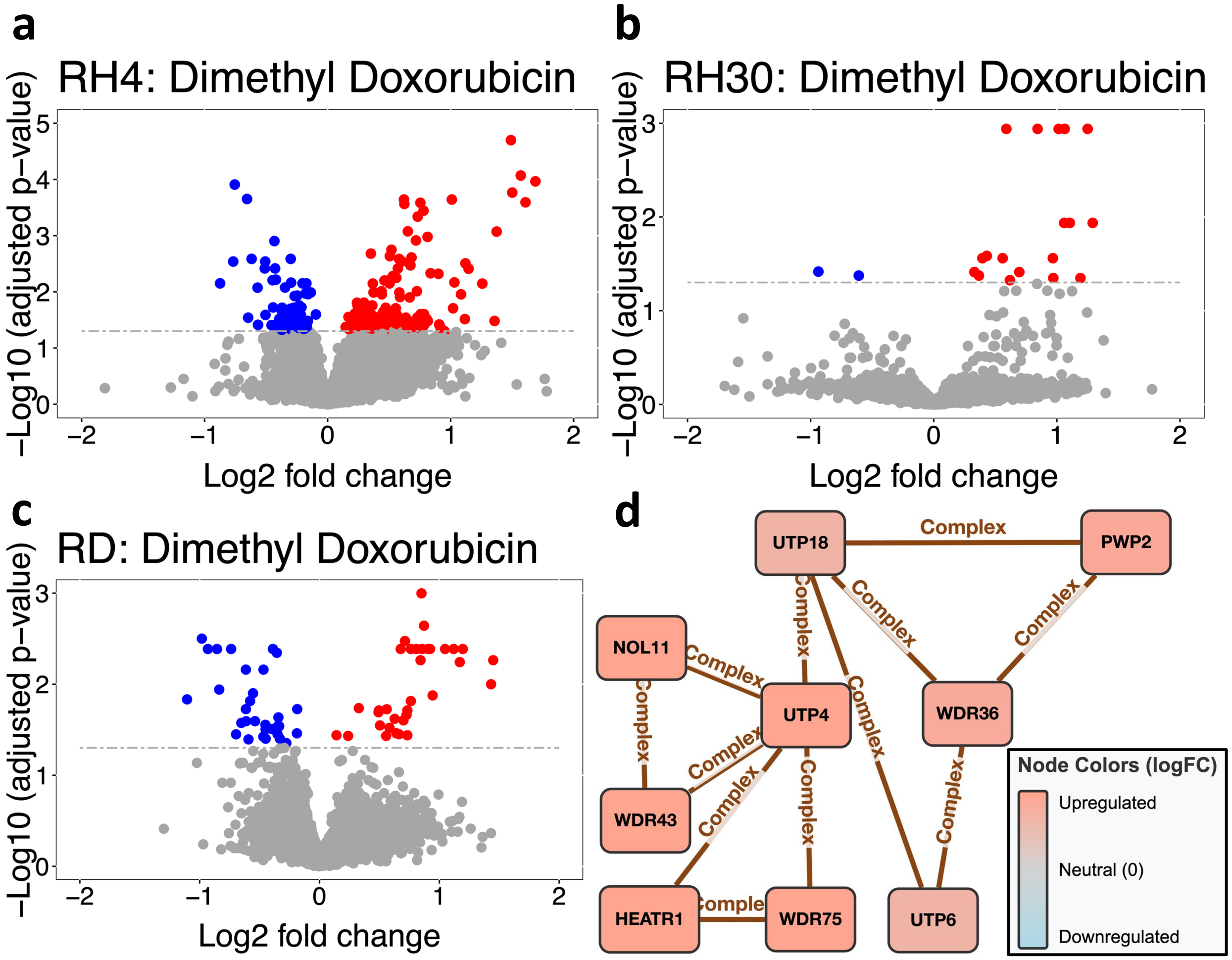

Figure S5: **RMS cell lines study: Volcano plots and corresponding subnetwork of common mechanisms across three cell lines.** a) Volcano plot of dimethyl doxorubicin compared to DMSO for the RH4 cell line, with significant proteins in blue and red. b) Same as (a) except for the RH30 cell line. c) Same as (a) except for the RD cell line. d) Subnetwork of common mechanisms across differential abundance analysis results of the RH4, RH30, and RD cell lines, color coded by direction of regulation and magnitude of median $\log_2$ fold change across cell lines.

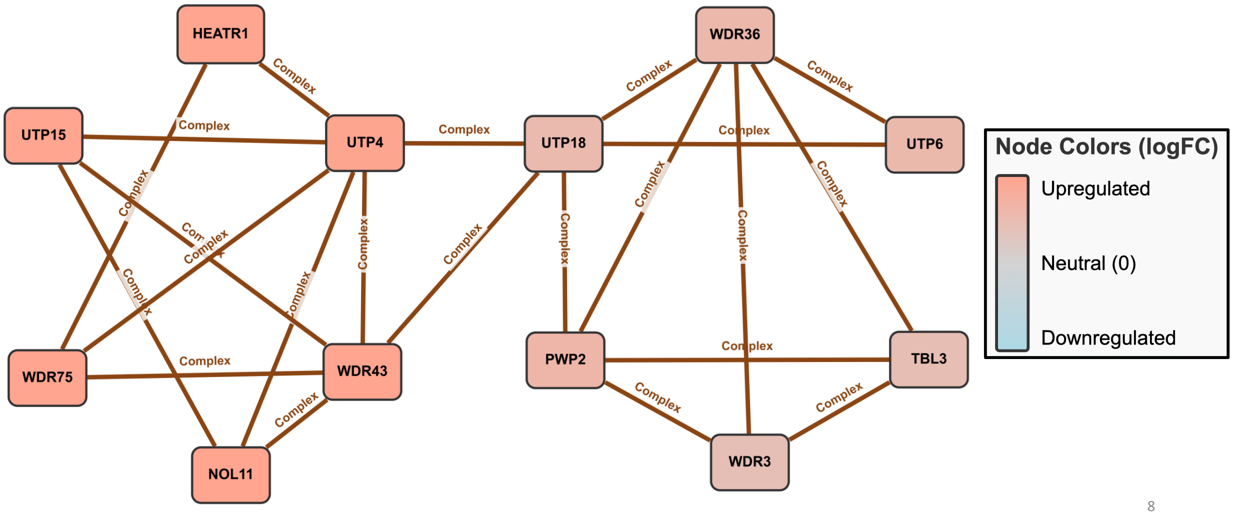

Figure S6: **RMS cell lines study: Protein subnetwork extracted with MSstatsBioNet with significant proteins and additional proteins of interest UTP15 and TBL3 given their involvement in UtpA and UtpB respectively.** On the left, UtpA protein complex consists of HEATR, UTP4, WDR43, NOL11, WDR75, UTP15. On the right, UtpB protein complex consists of UTP18, UTP6, TBL3, PWP2, WDR36, WDR3. Node color represents median $\log_2$ fold change magnitude across cell lines.

#### S3 IFN$\gamma$ stimulation and *LGALS1* knockdown study: supplementary materials

Table S5: **IFN$\gamma$ stimulation and *LGALS1* knockdown study:** Protein-level subnetwork search summary. Comparison type refers to whether the comparison answers a question regarding the effect of IFN$\gamma$ stimulation, *LGALS1* knockdown, or their interaction

| Comparison | Comparison Type | No. Differentially Abundant Phosphosites | No. of INDRA statements |
|---|---|---|---|
| *LGALS1* siRNA + vehicle vs non-targeting control + vehicle | *LGALS1* knockdown | 10 | 2 |
| *LGALS1* siRNA + IFN$\gamma$ vs non-targeting control + vehicle | Interaction effect | 4 | 0 |
| non-targeting control + IFN$\gamma$ vs non-targeting control + vehicle | IFN$\gamma$ stimulation | 5 | 6 |
| *LGALS1* siRNA + IFN$\gamma$ vs *LGALS1* siRNA + vehicle | IFN$\gamma$ stimulation | 1 | 0 |
| *LGALS1* siRNA + vehicle vs non-targeting control + IFN$\gamma$ | Interaction effect | 3 | 0 |
| *LGALS1* siRNA + IFN$\gamma$ vs non-targeting control + IFN$\gamma$ | *LGALS1* knockdown | 4 | 0 |

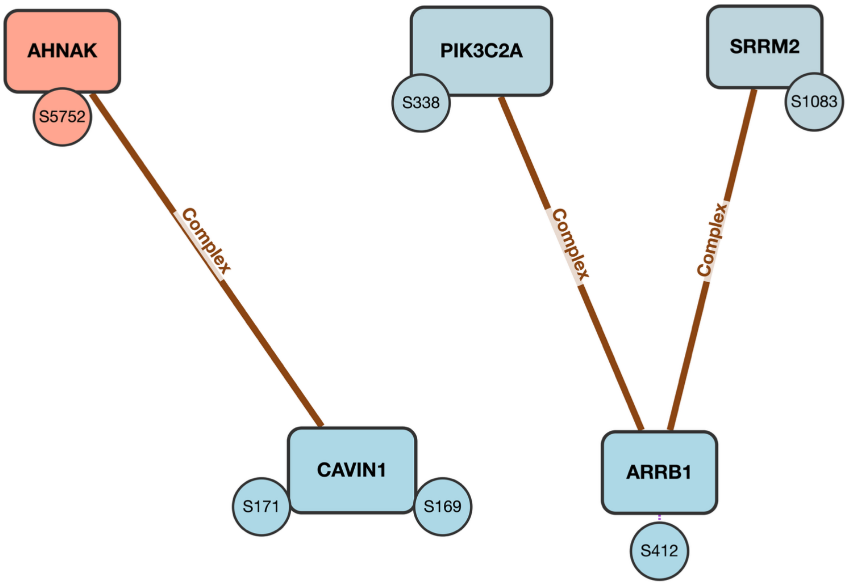

Figure S7: **IFN$\gamma$ stimulation and *LGALS1* knockdown study:** Subnetwork without PTM site specific relationships comparing non-targeting control + IFN$\gamma$ vs non-targeting control + vehicle (examining the effect of IFN$\gamma$ stimulation). Protein subnetwork with differentially abundant phosphosites, filtering by adj.pvalue $< 0.2$, INDRA evidence count $\geq 1$.

Table S6: **IFN$\gamma$ stimulation and *LGALS1* knockdown study:** Site-specific subnetwork search summary. Comparison type refers to whether the comparison answers a question regarding the effect of IFN$\gamma$ stimulation, *LGALS1* knockdown, or their interaction

| Comparison | Comparison Type | No. of Phosphosites Missing in One Condition | No. of INDRA statements |
|---|---|---|---|
| *LGALS1* siRNA + vehicle vs non-targeting control + vehicle | *LGALS1* knockdown | 496 | 9 |
| *LGALS1* siRNA + IFN$\gamma$ vs non-targeting control + vehicle | Interaction effect | 521 | 7 |
| non-targeting control + IFN$\gamma$ vs non-targeting control + vehicle | IFN$\gamma$ stimulation | 569 | 20 |
| *LGALS1* siRNA + IFN$\gamma$ vs *LGALS1* siRNA + vehicle | IFN$\gamma$ stimulation | 445 | 4 |
| *LGALS1* siRNA + vehicle vs non-targeting control + IFN$\gamma$ | Interaction effect | 553 | 15 |
| *LGALS1* siRNA + IFN$\gamma$ vs non-targeting control + IFN$\gamma$ | *LGALS1* knockdown | 580 | 17 |

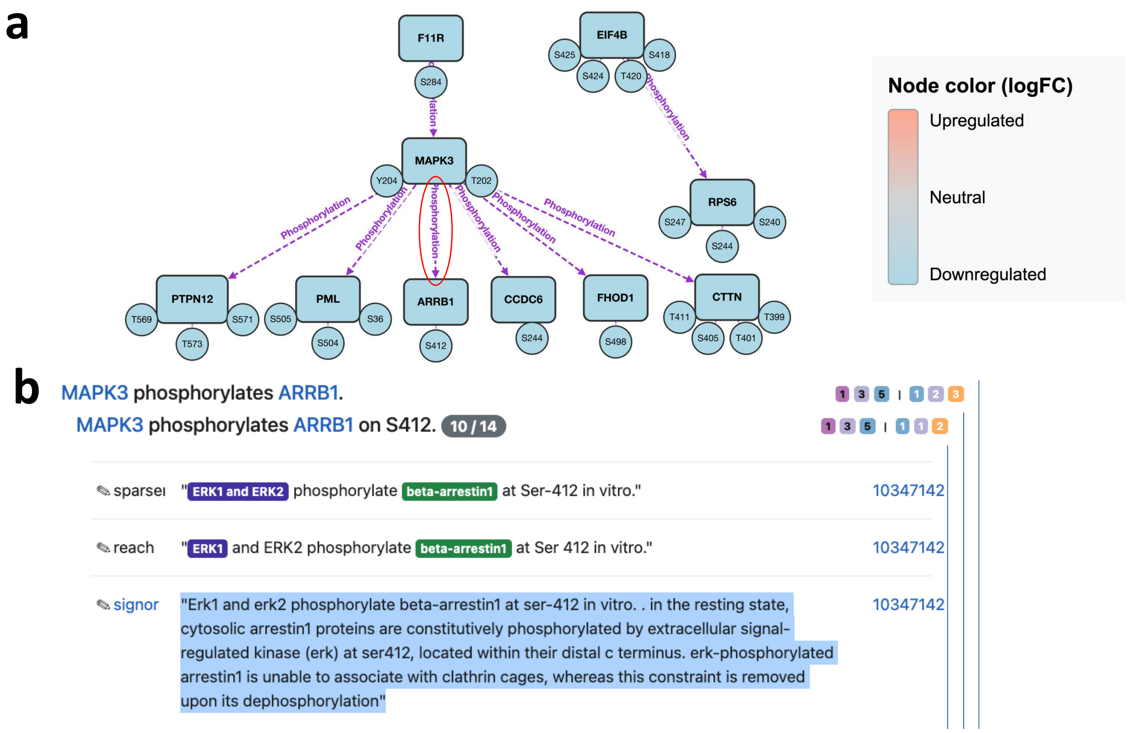

Figure S8: **IFN$\gamma$ stimulation and *LGALS1* knockdown study:** Subnetwork with PTM site specific relationships comparing non-targeting control + IFN$\gamma$ vs non-targeting control + vehicle (examining the effect of IFN$\gamma$ stimulation) a) Subnetwork with differentially abundant phosphosites, including phosphosites not measured under the non-targeting control + IFN$\gamma$ condition, filtered by PTM site specific relationships and down-regulated phosphosites. All of the phosphosites are not measured under the non-targeting control + IFN$\gamma$ condition but measured in all remaining conditions (with the exception of only ARRB1 at serine 412, which was measured in all conditions). b) INDRA web portal screenshot of the evidence text that MAPK3 phosphorylates ARRB1 at serine 412.

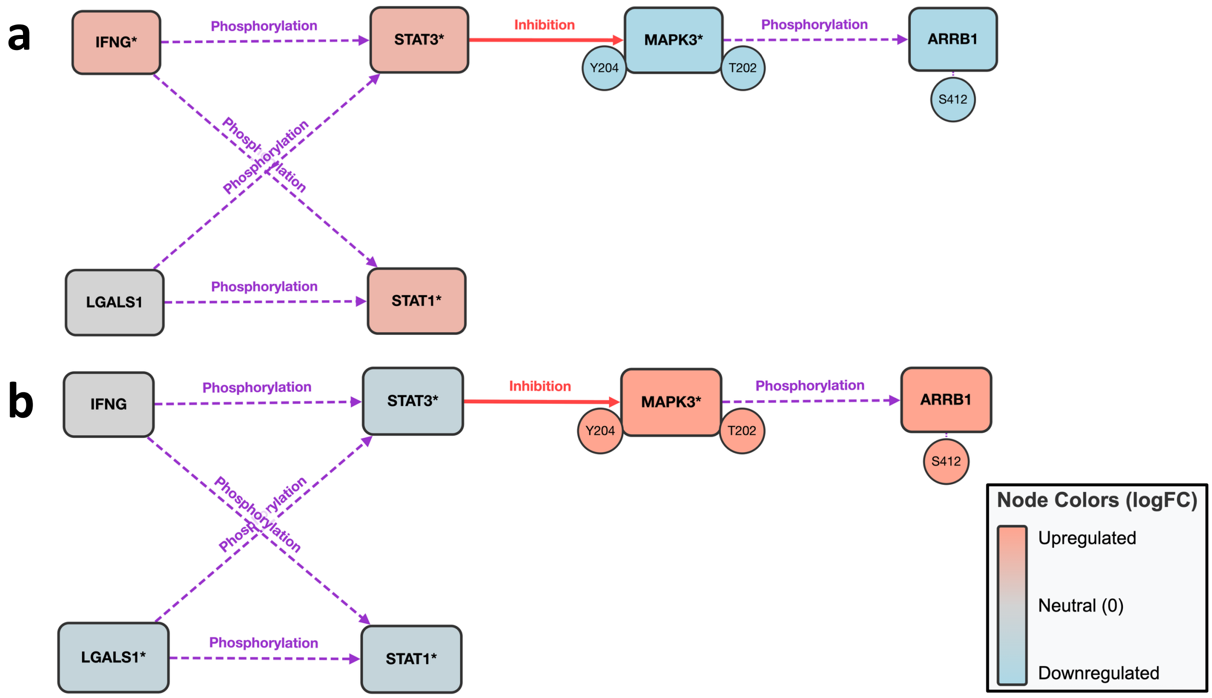

Figure S9: **IFN$\gamma$ stimulation and *LGALS1* knockdown study:** Subnetwork connecting MAPK3 to IFN$\gamma$ canonicial pathway. Phosphosites with their direction of regulation inferred based on assumptions are marked with an asterisk. a) Subnetwork with IFN$\gamma$ canonical pathway proteins and MAPK3 and ARRB1, with their direction of regulation derived from comparing non-targeting control + IFN$\gamma$ vs non-targeting control + vehicle (examining the effect of IFN$\gamma$ stimulation). b) Subnetwork with IFN$\gamma$ canonical pathway proteins and MAPK3 and ARRB1, with their direction of regulation derived from comparing *LGALS1* siRNA + IFN$\gamma$ vs non-targeting control + IFN$\gamma$.
